# A dynamic gradient architecture generates brain activity states

**DOI:** 10.1101/2020.08.12.248112

**Authors:** Jesse A. Brown, Alex J. Lee, Lorenzo Pasquini, William W. Seeley

## Abstract

A central goal of systems neuroscience is to determine the functional-anatomical basis of brain-wide activity dynamics. While brain activity patterns appear to be low-dimensional and guided by spatial gradients, the set of gradients remains provisional and their mode of interaction is unclear. Here we applied deep learning-based dimensionality reduction to task-free fMRI images to derive an intrinsic latent space of human brain activity. Each dimension represented a discrete, dynamically fluctuating spatial activity gradient. The principal dimension was a novel unipolar sensory-association gradient underlying the global signal. A small set of gradients appeared to underlie key functional connectomics phenomena. Different task activation patterns were generated by gradients adopting task-specific configurations. Dynamical systems modelling revealed that gradients interact via state-specific coupling parameters, allowing accurate forecasts and simulations of task-specific brain activity. Together, these findings indicate that a small set of dynamic, interacting gradients create the repertoire of possible brain activity states.

## Main Text

### Introduction

Functional connectivity is defined as synchronous activity within two or more brain regions over time. Functional connectivity patterns as revealed by functional MRI (fMRI) have advanced our understanding of the brain’s functional neuroanatomy in health (Yeo et al., 2011) and disease (Seeley et al., 2009). Despite this progress, the basis for functional connectivity remains unclear. Any candidate model must account for the diverse but constrained range of dynamic functional connectivity states that are observed both within and across individuals (Allen et al., 2014; Gratton et al., 2018; Pasquini et al., 2020; Vidaurre et al., 2017). Importantly, this limited flexibility of whole-brain activity suggests that a low-dimensional set of neuroanatomical systems may be involved (Glomb et al., 2019; Saggar et al., 2018; Shine et al., 2019). Recent work indicates that these dimensions reflect “gradients” of continuous variation in regional functional connectivity (Haak et al., 2018; Margulies et al., 2016; Zhang et al., 2019) and cytoarchitecure (Burt et al., 2018; Paquola et al., 2019; Wang, 2020), with clustered regions forming functionally connected networks. To date, however, no study has demonstrated how gradients and their dynamic interactions can collectively explain key observations in functional connectomics, including the brain-wide global signal (Fox et al., 2009), anti-correlation between large-scale networks (Fox et al., 2005), the presence of discrete functional modules and hub regions (Sporns and Betzel, 2016), correspondence with spatial patterns of gene expression (Richiardi et al., 2015), dynamic configuration into task-specific brain activity states (Barch et al., 2013), and stable functional connectivity patterns in individuals over time (Finn et al., 2015).

Here we apply deep-learning based dimensionality reduction to derive an intrinsic latent space of brain activity dynamics. Our approach explicitly focuses on instantaneous brain activity patterns and differs from conventional methods for deriving functional connectivity gradients (Hong et al., 2020). This approach yields a more comprehensive set of latent brain activity dimensions each associated with a spatial gradient. Most importantly, this includes a principal activity gradient that underlies the global signal of brain activity and that we believe is fully described here for the first time. We demonstrate that this expanded set of activity gradients represent intrinsic systems that are stable across individuals, multiple days of scanning, and task-free or task-engaged scanning conditions. In confirmatory analyses, we show that this set of latent dimensions and gradients capture essential properties of brain activity and functional connectivity including modularity and regional hub properties, correspondence with spatial gene expression patterns, and reconfiguration during specific mental tasks. Finally, we use dynamical systems analysis to reveal how this set of activity gradients can transiently couple into state-specific configurations, suggesting a novel mechanism the brain appears to use to dynamically generate diverse activity and functional connectivity states.

### Results

#### Low-dimensional brain activity latent space

We assessed latent brain activity dynamics in task-free and task-engaged fMRI scans by first applying a convolutional autoencoder, a deep learning tool for dimensionality reduction and the representation of spatial features from 2D or 3D images (Hinton and Salakhutdinov, 2006; Zeiler and Fergus, 2013). By design, convolutional neural networks can determine a more efficient data-driven spatial dimensionality reduction than techniques like independent component analysis (ICA) or principal component analysis (PCA) that are naïve to spatial structure in images (Calhoun et al., 2001; Beckmann and Smith, 2004). Autoencoders are also more computationally tractable than PCA for performing spatial dimensionality reduction on a large set of images. We therefore used a 3D convolutional autoencoder for spatial dimensionality reduction of 119,500 task-free fMRI images from a group of 100 healthy young control subjects in the Human Connectome Project (HCP; **Methods**). After autoencoder training, we obtained spatial embeddings for each image. We then ran subsequent PCA on the 119500 x 1080 embedding matrix to derive the latent space of functional brain activity with spatially and temporally orthogonal dimensions (**Figure 1A**). Component scores for each image were then linked to yield dynamic latent trajectories for each individual’s fMRI scan.

**Figure 1.**
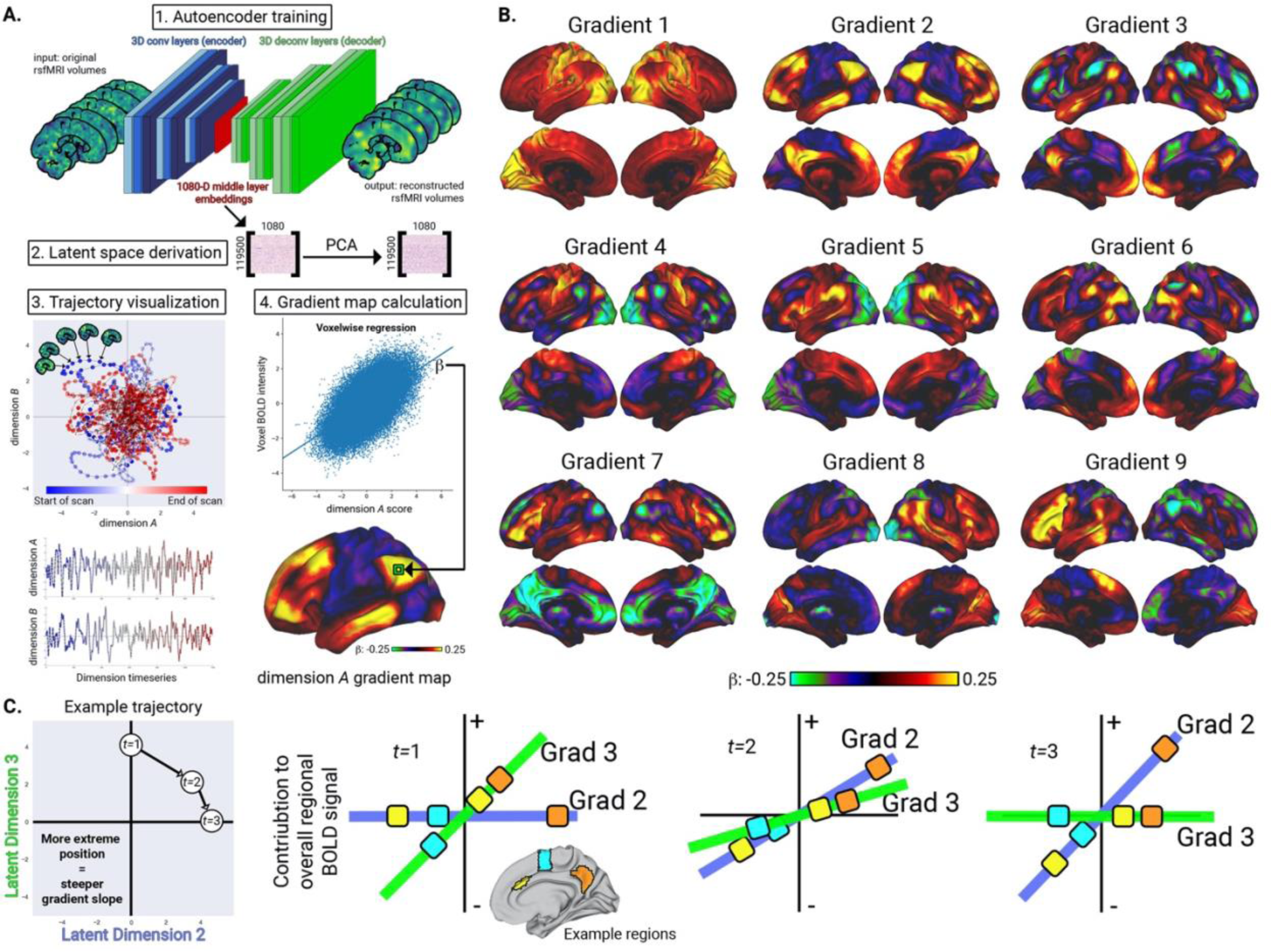
Latent space derivation, gradient maps, and temporal interpretation. A. The workflow for deriving individual latent trajectories and spatial activity gradient maps from task-free fMRI images. B. Activity gradient maps for the first nine latent space dimensions. The signs are arbitrary, representing only whether regions are correlated or anti-correlated with each other on that dimension. C. Illustration of relationship between latent space trajectories and regional BOLD activity. Across three successive timepoints, latent space positions on each dimension reflect the current slopes of the corresponding gradients. The resultant BOLD signal in each region depends on the region’s weight on each gradient, shown here for the anterior cingulate cortex (yellow), premotor cortex (cyan), and precuneus (orange).

In this latent space each dimension represented an independent spatial activity gradient and the entire space collectively encapsulated the range of possible brain activity states. To derive the spatial activity gradients associated with each latent dimension, we regressed each voxel’s BOLD activity against that latent dimension timeseries, inferring a voxel’s positive or negative beta weight on that dimension (**Figure 1A**). A voxel’s BOLD signal could be reconstructed by multiplying the voxel’s weight on a gradient by the gradient’s current slope and summing across all gradients (**Figure 1C, Movie S1,** and **Methods**). Throughout this study, we use the term “gradient” to refer to the spatial map associated with a latent dimension (analogous to loadings on a PCA component), the term “slope” to describe the steepness of the gradient at a specific timepoint (i.e. score on a PCA component), and the term “trajectory” in reference to a multi-dimensional set of gradient slope timeseries (i.e. a set of PCA score timeseries). When reconstructing each brain region’s timeseries, the first three dimensions explained 44.9% of BOLD activity variance across 273 cortical, subcortical, and cerebellar regions, while nine dimensions explained 57.9% **(Figure S1** and **Supplementary Information)**. This indicates that a low-dimensional latent space was able to explain a substantial proportion of the variation in BOLD activity. Our subsequent analysis focused on the first nine dimensions based on the criterion that diminishing additional variance was explained with subsequent dimensions (**Methods**).

The polarity of regional “positive” versus “negative” weights on a gradient was arbitrary, signifying only whether regions were correlated or anti-correlated with each other on that gradient (**Figure 1C**). These activity gradients demonstrated both similarities and key differences with functional connectivity gradients like the set described by Margulies and colleagues (referred to as FCG1-FCG5) (Margulies et al., 2016). The strongest spatial matches between each functional connectivity gradient and the activity gradients reported here were: FCG1/Gradient 2, r=0.86; FCG2/Gradient 5, r=0.76; FCG3/Gradient 3, r=0.88; FCG4/Gradient 7, r=0.57; and FCG5/Gradient 6, r=0.61. Overall, the activity gradients had strong correspondence with known functional connectivity gradients, with Gradient 1 being an important exception.

The spatial weights for Gradient 1 were positive across 99.6% of the gray matter (**Figure 1B**), albeit with topographically varied weights that were highest in the primary sensory, visual, and auditory cortices. The gradient slope timeseries associated with this dimension had a near-perfect correlation with the global gray matter signal (r=0.92; **Figure 2A**), a major influence on the estimated strength of functional connectivity (Fox et al., 2009). Gradient 1 was the only “unipolar” gradient, as no other gradient was more than 70% skewed towards positive- or negative-predominant weights brain-wide. We ensured that this unipolar component was not a statistical artifact by demeaning the data in two stages, both voxel-wise demeaning of BOLD activity before autoencoder inference and dimension-wise demeaning of spatial embeddings before PCA. Crucially, Gradient 1 could only be detected when performing dimensionality reduction on data with a temporal dimension. This approach differs from how spatial gradients are typically derived from fMRI data, by applying dimensionality reduction on the functional connectivity matrix. There is a straightforward reason why this unipolar gradient can be detected from the timeseries data but not from a functional connectivity matrix. PCA and related dimensionality reduction methods find components that maximize the variance in a measure across observations. When applied to timeseries data, PCA is capable of detecting a global unipolar spatial factor which causes all regions to have high or low activation at different timepoints. By contrast, in a functional connectivity matrix where the time dimension is collapsed, all regions would share a common pattern of connectivity reflecting the average slope of the unipolar Gradient 1. This lack of region-to-region variability in connectivity would render this factor invisible to the variance-maximizing PCA algorithm. This highlights the benefit of deriving spatial gradients from continuous BOLD activity data.

**Figure 2.**
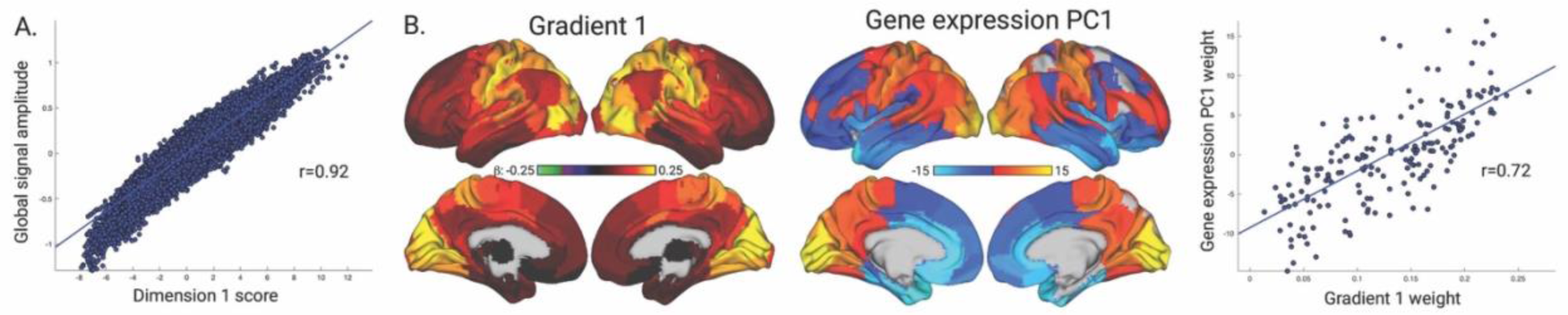
Gradient 1 underlies the global signal and corresponds to the principial spatial gene expression component. **A.** The correlation between the latent dimension 1 score, reflecting the gradient’s current slope, and the global gray matter signal. **B.** Spatial maps for Gradient 1 and the primary spatial component of genetic expression variability, along with their spatial correlation.

The spatial activity gradient for dimension 2 corresponded to the putative primary functional connectivity gradient (Margulies et al., 2016), a sensory-to-cognitive axis (**Figure 1B**) reflecting a functional spectrum from perception and action to more abstract cognitive function. The most positive weights were in areas of the default mode network and executive control network (**Figure S4** for overlap with seven canonical networks (Yeo et al., 2011)), while the most negative weights were in somatomotor, visual, and ventral attention networks. Dimension 3 resembled a task-positive (frontoparietal) to task-negative (default mode) gradient (**Figure 1B** and **Figure S4)**. The subsequent dimensions included strongly lateralized activity gradients (Gradients 8 and 9), oppositions between specific sensory modalities like the visual and somatomotor networks (Gradients 4 and 5), and differential involvement of sub-components of larger super-systems like the default mode network (Gradients 2, 3, and 7) (Andrews-Hanna et al., 2010). Critically, gradients were highly reproducible in an independent validation dataset with the first 12 dimensions appearing in the nearly the same sequence (all spatial correlation coefficients > r=0.5; **Table S1** and **Figure S4/Figure S5)**. Overall, we found that by performing dimensionality reduction on BOLD timeseries data, we could detect an expanded set of activity gradients from those previously reported, including the unipolar Gradient 1 which explained the most variance in brain activity. The consistency of the spatial gradients across individuals suggests that they reflect intrinsic anatomical systems of brain functional organization. We next evaluated the ability of this set of activity gradients to explain core phenomena in functional brain activity and connectivity.

#### Latent trajectories capture individual brain activity fingerprints

Importantly, individual subject latent trajectories (described by the gradient slope covariance matrix, **Methods**) exhibited high reliability on consecutive days of scanning, consistent with the finding of identifiable individual functional connectivity fingerprints (Finn et al., 2015). Six latent dimensions were required to correctly match a subject’s day 1 and day 2 scans for at least 50% of individuals, while 29 dimensions were required to identify the correct fingerprint for all 100 subjects (**Figure S6** and **Supplementary Information**). Thus, a low dimensional latent space of brain activity captures sufficient information to distinguish individuals, suggesting that reliable differences in gradient engagement may represent individual traits.

#### Basis for functional modularity and hubness

We next characterized how latent activity trajectories represent observed patterns of functional connectivity. Here we demonstrate how a latent trajectory can maximize functional connectivity (i.e. co-variation) between two example regions, the anterior cingulate cortex (ACC) and middle frontal gyrus (MFG), two regions of interest that belong to dissociable functional networks. We focused on Gradient 2 and 3, the first two bipolar gradients. We first determined which latent space trajectory direction was optimal for maximizing the ACC or MFG’s BOLD activation relative to all other regions (**Figure 3A**). The ACC’s activity was maximized by a mostly downward trajectory (**Figure 3A, top**). By contrast, the MFG’s activity was maximized by a more rightward trajectory (**Figure 3A, bottom**). The optimal trajectory angle to maximize co-variation between the ACC and MFG was from the top-left to the bottom-right of this 2D latent space, bisecting the trajectory angles that maximized activity in either the ACC or MFG. We confirmed that the individual subjects with maximal or minimal ACC-MFG functional connectivity during task-free fMRI had latent trajectories that were most or least aligned with the optimal co-variation angle (**Figure 3B; Methods** and **Supplementary Information)**. This illustrates the value of rendering brain activity as a latent trajectory, providing a read out for levels of regional activity or functional connectivity for any timespan, from a single timepoint to a full scan.

**Figure 3.**
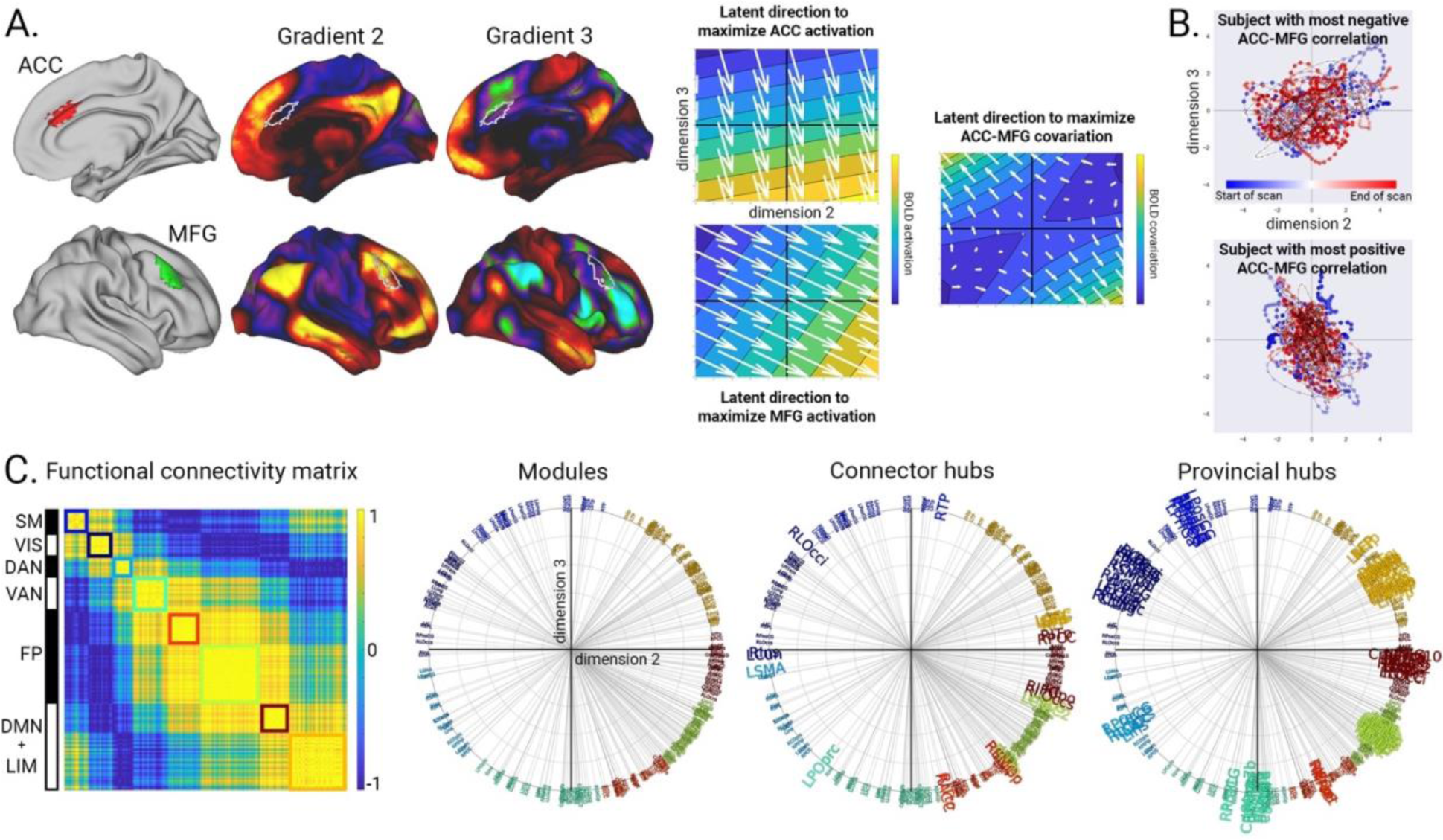
Functional modularity and hubness reflect non-uniform spacing of regions along gradients. **A.** Latent trajectory directions that maximize BOLD activation for the ACC, MFG, or the covariation between them. **B.** Latent trajectories during task-free scans for subjects with the most negative (top) or most positive (bottom) ACC-MFG correlation. **C.** Left: group-average functional connectivity matrix, based on region timeseries reconstructed only from latent dimensions 2 and 3. The eight modules detected in this connectivity matrix are highlighted along the diagonal along with the best-corresponding canonical network. Middle/Right: polar plots showing the best latent trajectory direction for maximizing activation in each region, which span the full 360° of the 2D latent space. Region colors are based on the modules they belong to, revealing that the bunching of regions with similar angles naturally reflects the modularity. Regions identified as the top connector hubs (larger font, middle) are the most distant (i.e. have the largest angles) from their neighboring regions in this 2D space, while provincial hubs (larger font, right) are the closest to their neighbors. SM: somatomotor, VIS: visual, DAN: dorsal attention network, VAN: ventral attention network, FP: frontoparietal, DMN: default mode network, LIM: limbic. See deposited data for full list of region abbreviations.

The existence of continuous spatial activity gradients may appear to be at odds with the presence of discrete modular brain networks, a major principle of brain functional organization (Sporns and Betzel, 2016). We therefore attempted to reconcile the gradient and modular perspectives by testing the hypothesis that non-uniform spacing of regions along a gradient would recapitulate modular boundaries. By assessing the optimal trajectory direction for maximizing BOLD activation in every brain region, we discovered that the trajectory angles fully spanned the 360° of the 2D latent space (**Figure 3C**). Regions tended to cluster with their contralateral homologues and other regions belonging to the same functional connectivity network, while regions that were diametrically opposed belonged to canonically anti-correlated networks. Based on this observation, we expected that functional connectivity modules derived from the functional connectome would correspond well with different angular ranges in latent space. We found that region module membership from the functional connectome corresponded exactly to the sequence of regions as grouped by optimal activation angle (**Figure 3C**).

Consequently, provincial hubs and connector hubs were found to have characteristic orientations. Provincial hubs had significantly smaller angles to their most strongly connected neighbors than non-hubs (provincial hubs: mean=2.8°±2.3°; non-hubs: mean=4.0°±3.7°; F=80.45, p=5.72×10^-19^; **Figure S8**), which in turn had smaller angles to their neighbors than connector hubs (connector hubs: mean=4.8°±3.9°; F=9.94, p=0.001). This relationship held true when considering a higher dimensional latent space with a larger number of gradients (**Supplementary Information**). Thus, the presence of modularity and hub regions appears to be consistent with the non-uniform spacing of regions along gradients.

#### Correspondence with spatial gene expression patterns

The spatial gradients described here have a striking congruence with morphogen gradients that guide regional differentiation and connectivity during brain development (O’Leary et al., 2007). This motivated a systematic assessment by comparing each activity gradient’s spatial similarity with genetic expression maps using the 15,655 genes across 196 cortical regions from the Allen Human Brain Atlas (**Methods**). While numerous relationships between structural or functional gradients and spatial gene expression patterns have been established (Burt et al., 2018; Vogel et al., 2020; Shafiei et al., 2020; Huntenburg et al., 2021), we sought to determine if the activity gradients defined in this study had stronger correspondence to gene expression patterns than to functional gradients derived by other methods. Across the first nine gradients, 4089 genes showed significant spatial correlations with at least one gradient in both the discovery and validation datasets, with correlation coefficients ranging between r=0.37-0.81 (surviving Bonferroni corrected threshold p < 3.55 x 10^-7^ in both the discovery and validation datasets). The most striking correspondences were with gradient 1, for which 3572 genes were significantly correlated and which explained up to 64% of the variance in regional gradient weight. A gene ontology enrichment analysis based on the full set of significantly correlated genes found associations including “ion gated channel activity”, “neuron projection”, “synapse”, and “anatomical structure development” (top-ranked terms in **Table S2**, all terms in deposited data). The most significant positive relationships between gene expression and gradient 1 were *SEMA7A* (discovery/validation mean r=0.78), *SCN1B* (r=0.76), *LAG3* (r=0.76), *ACAN* (r=0.76), *ASB13* (r=0.76), *SV2C* (r=0.76), *ANK1* (r=0.76), *and GPAT3* (r=0.76), while the most negative relationships were *KCNG1* (r=-0.81), *ASCL2* (r=-0.78), *ANKRD6* (r=-0.76), *PRKCD* (r=-0.75), and *RSPH9* (r=-0.75). Because such a large number of genes exhibited strong positive or negative spatial correlations with gradient 1, we tested for a potential link to the primary spatial component of genetic expression variability, which is known to stratify sensory and association areas (Burt et al., 2018). There was a significant spatial correlation between Gradient 1 and the principal spatial component of gene expression (r=0.72; **Figure 2B**), for which each of the aforementioned individual genes were strongly loaded (all loading absolute Z scores > 2.8). This correlation coefficient is stronger than any previously reported correlation between functional gradients and human spatial gene expression patterns that we are aware of in the literature.

This indicated that the predominant sources of variability in BOLD activity and spatial gene expression are strongly linked.

Substantially fewer genes were significantly correlated with the remaining gradients (**Supplementary Information** and **Figure S9**). Gradient 2 had 48 significantly correlated genes while interestingly, gradient 3 had no significantly correlated genes (discovery minimum p=2.51 x 10^-6^, validation minimum p=9.06 x 10^-11^) despite explaining a substantial portion of BOLD variance, having a replicated spatial pattern in the validation dataset, and showing spatial correspondence to a previously described functional connectivity gradient (**Supplementary Information**). The lack of genetic correspondence for this gradient may be due to the stringent criterion for statistical significance or greater individual variability. Among the strongest of the other gene/gradient relationships were *CARTPT* on gradient 2 (r=0.55) and *CDH6*, *CDH13,* and *FABP6* on gradient 5 (r=0.60/0.58/0.60). Genes associated with activity gradients were recurrently linked to functional and structural processes likely to influence brain-wide activity patterns including anatomical morphogenesis, excitation/inhibition balance, and thalamocortical connectivity.

#### Gradient-based task-induced brain activity patterns

We next evaluated the possibility that activity gradients detected during the task-free state are spatially stable, intrinsic systems that can dynamically adopt specific configurations to create task-specific brain activity states. The key novel aspect of our approach was projecting task fMRI data into the task-free defined latent space, based on our hypothesis that the dimensions and associated gradients are intrinsic to the brain and will remain fixed, while the shape of the trajectories will change in task-specific fashion. We thus projected HCP task fMRI data from validation dataset subjects into the task-free latent space defined from discovery dataset subjects, then derived gradient slope timeseries for each condition in each task. We focused on four diverse cognitive tasks known to elicit distinct brain activity patterns – working memory (2-back vs. 0-back), motor movement (finger/toe/tongue movement vs. visual fixation), language comprehension (auditory story vs. math questions), and emotion processing (face emotion recognition vs. shape recognition). In each task, we found selective gradient slope differences between conditions (abs(t) > 3.17, p < 0.005) that combined to produce greater activation in areas consistent with previously reported task-activity patterns (**Figure 4**) (Barch et al., 2013). We verified the plausibility of the task-specific gradient-based activation patterns by measuring their spatial correlation with conventional voxel-wise GLM-based task activation maps. Gradient-based maps had strong correlation (r > 0.5) with the GLM-based maps when including between 3 to 10 gradients, rapidly increasing in correspondence before plateauing after ∼20 gradients and reaching a maximum of r=0.76-0.91. The rapid increase and subsequent plateau of correlation suggests that a low-dimensional set of activity gradients are sufficient to generate diverse task-activation patterns. Furthermore, this finding is evidence that the latent dimensions derived from the task-free state are intrinsic, and that what changes during a task is the engagement level of each gradient.

**Figure 4.**
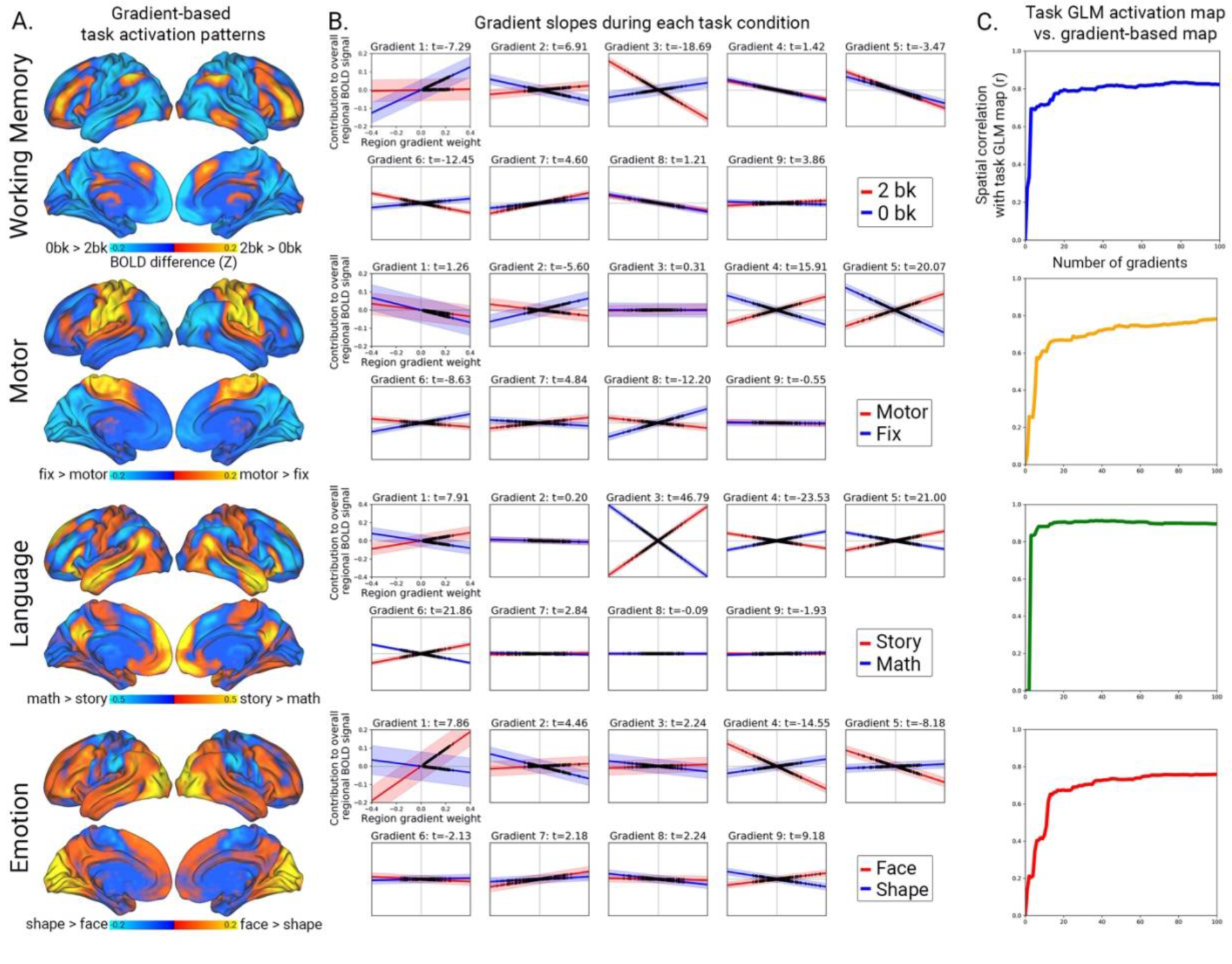
Specific gradient slopes produce task-specific activity patterns. **A.** Reconstructed BOLD activity differences between the active and baseline conditions in the working memory, motor, language, and emotion tasks, based on gradient slope differences of the first 9 latent dimensions after projecting task fMRI data into the task-free defined latent space. **B.** Gradient slope differences between the contrasting task conditions, where the line slope represents the across-subject average and the shaded band represents the 95% confidence interval. Black tick marks denote each region’s weight on each gradient. **C.** Spatial correlation between the gradient-based task activation maps from panel A and the task activation maps derived with a conventional general linear model (GLM).

#### Predicting state-specific dynamic trajectories of brain activity

The engagement of activity gradients at specific levels during different tasks raised the question of how gradients with perpetual dynamics can organize into distinct global states. This motivated our use of a dynamical systems model to infer how gradients influence each other in an ongoing fashion. Based on our observation that latent trajectories exhibited continuity and momentum, we chose to model the gradient slope timeseries with differential equations describing the continuous influence of each activity gradient on one another (Breakspear, 2017). We used a data-driven strategy to estimate the “coupling parameters” between gradients (Brunton et al., 2016). We first focused on the task-free fMRI data. For each fMRI timepoint, we measured the gradient slope (*g*), the first derivative of the slope (*g*’, the velocity of the change in slope), and the second derivative of the slope (*g*’’, the acceleration of the change in slope) (**Figure 5A**). We then used linear regression for each of the first nine gradients to estimate gradient slope acceleration as a function of all gradients’ slopes and slope velocities. We found that a gradient’s slope acceleration *g*’’ primarily had a strong negative relationship with its own slope *g* (mean b=-0.06±0.007, mean t=-238.6±12.8), as is characteristic of an oscillating signal. Gradients also had selective relationships with the slope velocity *g’* of the other eight gradients (b absolute range=0.0001-0.10), t absolute range=0.13-37.09) (**Figure S10**). This demonstrated that a given gradient’s current activity level is dependent on changes in both its own activity and the activity of specific other gradients.

**Figure 5.**
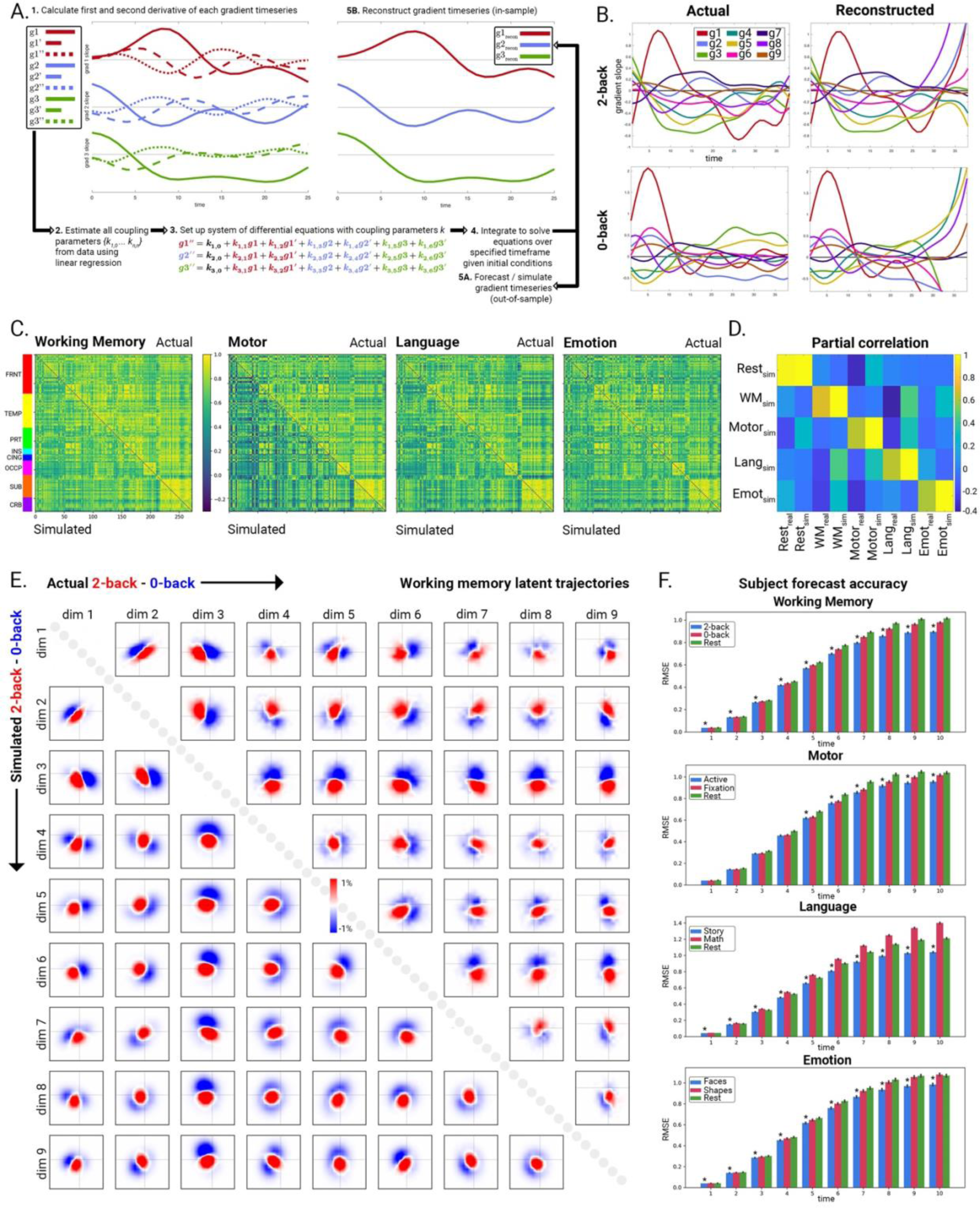
Distinct gradient coupling patterns produce state-specific functional activity and connectivity patterns. **A.** Gradient forecasting schematic. For each gradient, a linear model estimated the gradient’s acceleration as a function of all gradients’ slopes and first derivatives. The regression-derived coupling parameters were used to specify the system of coupled differential equations, which were then solved using initial conditions and coupling parameters that either come from in-sample data (“reconstructions”) or out of sample data (“forecasts”). **B.** Differential equation-based reconstructions of gradient timeseries during the 2-back and 0-back conditions of the working memory task, based on trial-locked group-average data. The reconstructions accurately match the actual data up until *t*=25 timepoints (18 seconds), after which they become unstable. **C.** Functional connectivity matrices from the active condition for each task, comparing actual data region pairwise correlations (upper triangle) with simulated data using state-specific coupling parameters (lower triangle). FRNT: frontal, TEMP: temporal, PRT: parietal, INS: insula, CING: cingulate, OCCP: occipital, SUB: subcortical, CRB: cerebellar. **D.** Partial correlations between each simulated functional connectivity matrix and the real/simulated functional connectivity matrices. **E.** Actual and simulated latent trajectories during working memory. Latent space dwell time differences are shown for the first nine latent dimensions, with actual working memory task trajectories on the matrix upper triangle and simulated trajectories on the lower triangle. Latent space locations where trajectories dwell more during the 2-back than the 0-back condition are shown in red and vice versa in blue. **F.** Task fMRI forecast accuracy. In each task, coupling parameters from condition of interest (blue; *: significantly lower than red, FDR corrected) yielded more accurate forecasts (root mean squared error ± standard error of the mean) than task-free state parameters (green), or opposite task condition coupling parameters (red).

In the next step, we used these coupling parameters to model nine-dimensional trajectories as a system of second-order differential equations (**Figure 5A** and **Methods**). We measured the accuracy of these differential equations by using them to generate forecasts of task-free state latent trajectories from an initial starting condition. Specifically, starting with initial slope and velocity of the nine gradients at a given timepoint *t* (randomly selected from the actual task-free fMRI data), we predicted the gradient slope timeseries for 10 future timepoints *t*+1 to *t*+10. Critically, we used coupling parameters derived from the discovery dataset to make forecasts in the validation dataset. As a benchmark for this forecast accuracy, we compared these forecasts to alternative forecasts generated by a first-order autoregressive model, a standard timeseries forecasting procedure which makes predictions based on the previous timepoint by leveraging the autocorrelation in the signal (Liégeois et al., 2017). The differential equation-based forecasts of the gradient slopes for gradients 1-9 at the subsequent fMRI timepoints explained significantly more variance than the autoregressive model (all p < 0.001): *t*=2, 99.8%/94.0% (Z=385.0); *t*=3, 96.9%/78.2% (Z=244.5); *t*=4, 87.3%/58.1% (Z=162.5); *t*=5, 69.2%/38.8% (Z=105.2); *t*=6, 46.1%/23.9% (Z=60.7); *t*=7, 24.7%/13.9% (Z=26.9); *t*=8, 10.3%/8.0% (Z=5.5). Thus, our model’s short-term forecasts outperformed a conventional timeseries forecasting technique. This provides evidence that a major determinant of near-term future brain activity patterns is the dynamic interaction of spatial activity gradients guided by specific coupling parameters.

Building on this finding, we hypothesized that specific gradient coupling modes would be required to induce task-specific activation patterns. We expected that forecasts of task-specific fMRI activity timeseries generated with the corresponding task-specific coupling parameters would be more accurate than forecasts using alternative coupling parameters, either from other tasks or the task-free state. We tested this hypothesis in three ways: task-specific gradient slope timeseries reconstruction, simulation, and forecasting (**Figure 5A**).

Reconstructions illustrated that the differential equations accurately modeled the within- and between-gradient factors that guide the moment-to-moment evolution of gradient changes for each task over a time horizon of ∼20 timepoints (∼14 seconds; **Figure 5B** and **Supplementary Information**). Next, we performed simulations to test whether differential equations with state-specific coupling parameters could generate timeseries that mimicked the brain-wide temporal dynamics in the actual task fMRI data that result in task-specific functional connectivity patterns (**Methods**). For each task, the simulated functional connectivity matrix generated from state-matched coupling parameters was always most similar to the true functional connectivity matrix from the matched task condition (cross-validated correlations: working memory, actual vs. simulated functional connectivity matrix r=0.95, Z=12.09; motor r=0.94, Z=5.33; language r=0.93, Z=7.83; emotion r=0.95, Z=21.45; all Z-associated p < 0.001; **Figure 5C** for real and simulated functional connectivity matrices; **Figure 5D** for partial correlations between simulated matrices and actual matrices; **Table S3** and **Supplementary Information)**. To illustrate the influence of the coupling parameters on latent trajectories, we compared latent space dwell time differences for the simulated state-specific latent trajectories to the true latent trajectories in the working memory task (**Figure 5E**). The shapes and locations of the 2D trajectories were consistent across multiple dimensions, indicating that the simulations can replicate higher-order geometrical properties of the multi-dimensional data manifold.

Finally, we performed timeseries forecasting to verify that state-matched coupling parameters could accurately extrapolate task-specific activity trajectories for unseen subjects. With parameters derived from the discovery dataset, the forecasts for all four tasks in the validation dataset were always significantly more accurate when based on the state-matched parameters than when using parameters from the baseline condition or from task-free state (working memory, for timepoints *t*+1 to *t*+10, all t-statistics versus baseline condition > 3, all FDR-corrected p < 0.01; motor, timepoints *t*+5 to *t*+10; language, timepoints *t*+1 to *t*+10; emotion, timepoints *t*+1 to *t*+10; **Figure 5F** and **Supplementary Information)**. Collectively, this set of experiments showed that differential equations with state-specific coupling parameters effectively captured the most salient aspects of task-specific brain activity including the dynamics of the task activity timeseries and the functional connectivity patterns.

### Discussion

This study makes three primary contributions to understanding the functional-anatomical basis of low-dimensional brain activity. First, we performed spatial dimensionality reduction on a large fMRI timeseries dataset using a convolutional autoencoder and subsequent PCA. By focusing the dimensionality reduction on timeseries data rather than a static functional connectivity matrix, we were able to derive a more encompassing set of latent brain activity dimensions than has previously been described. Most importantly, this set included a novel primary dimension of brain activity reflecting a unipolar spatial gradient most strongly weighted in unimodal cortical areas. This gradient explained the most variance in BOLD activity, appeared to underlie the global signal, and had a stronger correlation with the principal spatial component of gene expression than any previously described functional gradient. Our second contribution was to demonstrate that these latent dimensions and gradients appear to be intrinsic. We tested this by defining the latent space from task-free fMRI data, projecting task fMRI data into that latent space, and finding that different task activation patterns were generated by intrinsic gradients steepening or flattening into task-specific configurations. Our third contribution was to model gradient interactions as a dynamical system. We showed that coupling parameters determine the level of influence that gradients have on one another, and that a dynamical model with state-specific coupling parameters can yield accurate forecasts and simulations of functional connectivity during different tasks.

We performed several analyses to confirm that this expanded set of latent dimensions and gradients are consistent with core phenomena in functional connectomics. First, the gradient spatial patterns and ordering in the discovery cohort were reproduced in the validation cohort, providing evidence that this functional anatomy is stable across individuals. Second, the activity gradients we describe here had specific correspondences with known functional connectivity gradients (Margulies et al., 2016). Third, individual subjects had identifiable latent trajectories on consecutive days, indicating that functional connectivity fingerprints were apparent in the low-dimensional latent space (Finn et al., 2015). Fourth, regions were spaced non-uniformly along gradients, reflecting functional modularity and the presence of hub regions. Fifth, gradients had strong spatial correlations with spatial gene expression patterns.

Collectively, these novel and confirmatory analyses establish that the brain activity latent space captures the core properties of functional brain activity and connectivity.

The primary gradient had a unipolar activation pattern reflecting the global signal of brain activity. The global signal is a source of ongoing controversy in fMRI literature, appearing to have a neuronal basis (Turchi et al., 2018), relating to individual differences in behavior (Uddin, 2020), but also associating with respiration or head motion (Chang and Glover, 2009; Power et al., 2014). We used ICA-FIX denoised HCP fMRI data to minimize the impact of non-neural signals including respiration (Power et al., 2017; Glasser et al., 2019). We found that this gradient had strongest involvement of unimodal visual, somatomotor, and auditory areas, consistent with previous reports that the global signal has a heterogenous spatial topography (Liu et al., 2018b; Li et al., 2019; Orban et al., 2020). The unipolar nature of Gradient 1 is a likely reason why it has not previously been detected with conventional gradient discovery methods (Vos de Wael et al., 2020). Gradient 1 acts like a rising and falling tide, driving positive correlation between all areas. In this case, only methods which look for latent dimensions of variability in activity will detect this gradient, unlike methods which look for latent dimensions of functional connectivity. Similarly, recent work has shown that when the global signal is low or has been regressed out, the most prominent residual spatiotemporal pattern involves anti-correlation between the default mode and task positive networks (Yousefi et al., 2018; Yousefi and Keilholz, 2020), strongly resembling Gradient 2 in the current study. One practical implication of a latent “global signal” dimension existing is that cleanly isolating the global signal may be more effective using PCA or temporal ICA (Glasser et al., 2018) rather than conventional global signal regression.

Gradient 1 had a striking spatial correlation with the principal spatial component of cortical gene expression. This correlation is the strongest reported relationship we are aware of between a functional gradient and a spatial gene expression pattern, supporting its biological plausibility. The principal genetic expression component is known to have a strong relationship with the cortical myelination pattern that demarcates the borders between sensory and association areas (Burt et al., 2018). In the current study, the genes most positively associated with Gradient 1 included *SEMA7A* and *SCN1B*, which have known roles in excitation/inhibition balance and seizure disorders (Brackenbury et al., 2013; Carcea et al., 2014). Collectively, the primary gradient appears to represent a convergent system of functional, structural, and genetic variation in the brain which may modulate global neuronal excitation/inhibition balance (Wang, 2020).

Gradient 2 strongly resembled the previously described principal macroscale gradient of functional connectivity (Margulies et al., 2016). This gradient defines a sensory-to-cognitive axis with the default mode network at one extreme and somatomotor and visual areas at the other. Balanced anticorrelation between networks is a central aspect of brain functional connectivity (Fox et al., 2005) and could plausibly be instantiated by placing brain areas at opposing ends of a single dynamic gradient. What circuit or systems-level mechanism might drive the ongoing fluctuations of these global, bipolar gradients? One compelling possibility is the reciprocal inhibitory connections in the thalamic reticular nucleus, which excite one thalamic nucleus while inhibiting an opposing cellular cluster (Crabtree, 2018). This motif is essential for switching between attending to visual or auditory stimuli (Schmitt et al., 2017) and may enable thalamic coordination of widespread cortical functional connectivity (Hwang et al., 2017; Buckner and DiNicola, 2019). We found that multiple gradients had correlations with genes such as *SEMA7A* and *CDH6* that demarcate specific thalamic nuclei and are also expressed in cortical layers where thalamocortical axons terminate (Bibollet-Bahena et al., 2017). Proof of a causal link between microscale thalamic electrophysiology and macroscale BOLD activity gradient fluctuations may require an optogenetic fMRI approach (Liu et al., 2015).

One consequence of functional anticorrelation between systems is modularity, which the brain uses to perform segregated cognitive processing (Sporns and Betzel, 2016). Here we find that the presence of modular boundaries, provincial hubs, and connector hubs reflects the non-uniform spacing of regions along each gradient. This principle of non-uniform spacing or “clumpiness” is known to influence the distinctness of regional boundaries (Tian and Zalesky, 2018). Our observation that gradients had significant spatial correlations with gene expression maps suggests that a non-uniform distribution of gene expression along gradients is also likely, with sections of a gradient expressing either clustered or transitioning gene expression profiles. Canonical functional connectivity networks have similarly been shown to exhibit discrete spatial gene expression patterns (Bertolero et al., 2019; Richiardi et al., 2015). Morphogen gradients during brain development may provide the scaffold for the emergence of multiple distributed and overlapping activation gradients (O’Leary et al., 2007).

We found that task-specific activation patterns resulted from specific levels of gradient engagement. This supports the hypothesis that the gradients are spatially fixed, intrinsic systems. If this is true, then the primary means for the brain to occupy different states is for gradients to steepen or flatten different amounts. We observed that latent trajectories reflecting gradient slopes are bounded during different task conditions, as demonstrated for the working memory 2-back and 0-back conditions (**Figure 5E**). This builds on previous work showing that distinct low-dimensional trajectories underlie task-specific brain activity states (Shine et al., 2019). A key consequence of a small set of spatially fixed, intrinsic gradients underlying brain activity patterns is that the range of possible functional network configurations is limited. This is in line with studies showing that brain-wide functional connectivity patterns only modestly reconfigure between task-free and task-engaged states (Cole et al., 2014; Gratton et al., 2018).

Our dynamical systems model showed that distinct brain activity states can be reached when gradients interact following state-specific coupling parameters. As we illustrated for the working memory task, the shape of the latent brain activity manifold is dictated by the gradient coupling parameters. The shape of this manifold will determine the coactivation patterns occurring during that state (Liu et al., 2018a), the probability of transition between different states (Vidaurre et al., 2017), and the temporal sequence of activity flow (Cole et al., 2016). One critical aspect of our modeling approach is that while the PCA-derived gradient timeseries are temporally orthogonal, this does not imply that gradients cannot causally influence each other.

Ordinary differential equations can accurately capture non-linear relationships between orthogonal variables in a low-dimensional system (Brunton et al., 2016). Our discovery of specific coupling parameters generating unique functional connectivity states is evidence that the gradients do not function independently, but instead exert distinct influence on one another. The amount of push and pull between gradients may be calibrated by neuromodulation (Shine, 2019) or other mechanisms of gain control (Buzsáki, 2019). We observed a strong tendency for a transiently steep gradient slope to subsequently flatten out, suggesting that gradient engagement and maintenance may be energetically costly. A tendency towards relaxation in parallel with mutual influence between gradients may lead to a “frustrated equilibrium” that perpetuates dynamic activity (Gollo and Breakspear, 2014). Future work can seek to understand how gradient coupling evolves on short timescales and is altered by neurological conditions or modified by feedback over extended timescales.

### Materials and Methods

#### Subjects and data

200 subjects were randomly selected from the Human Connectome Project (HCP) 1200 Subjects Data Release with available resting (task-free) and task fMRI data from a 3T MRI scanner (https://db.humanconnectome.org/data/projects/HCP_1200). Informed consent was obtained for each individual by the HCP consortium. The HCP complied with all relevant ethical regulations. This study agreed to the Open Access Data Use Terms (https://www.humanconnectome.org/study/hcp-young-adult/document/wu-minn-hcp-consortium-open-access-data-use-terms) and was exempt from the UCSF IRB because investigators could not readily ascertain the identities of the individuals to whom the data belonged. Task-free state scans were 14.4 minutes long with a repetition time (TR) of 720 ms, resulting in 1200 fMRI volumes per scan. We divided these subjects into 100-subject discovery and validation datasets (56 female/44 male in the discovery dataset, mean age=28.9 years; 50 female/50 male in the validation dataset, mean age= 28.6 years). For task-free state fMRI data, we used the left-right phase encoded, minimally preprocessed scans with motion correction and FIX-ICA denoising. fMRI volumes were temporally concatenated for all subjects in the discovery or validation datasets (119500 volumes each) and each voxel’s timeseries was then standardized to have a mean of zero and a standard deviation of one. More detailed scanning parameters and preprocessing procedures have been described in detail elsewhere (Glasser et al., 2013). We used FSL (https://fsl.fmrib.ox.ac.uk/fsl/fslwiki/) and AFNI (https://afni.nimh.nih.gov/) for subsequent fMRI preprocessing. The first 5 volumes for each fMRI scan were dropped to allow scanner stabilization. Scans were bandpass filtered in the 0.008-0.15Hz frequency range and subsequently normalized in each voxel across time to have zero mean and unit variance.

#### Deep learning

All preprocessed fMRI volumes in this study were downsampled to 48x48x48 cubic images with 4 mm^3^ isotropic resolution, the maximal resolution possible due to GPU memory constraints during deep neural network training. We used a three-dimensional deep convolutional autoencoder (3D-DCA) to learn a dimensionality reduction of the 3D fMRI volumes. The 3D-DCA architecture we used was: 64 filter convolutional layer with a 3 voxel-by-3 voxel kernel and a stride of 2 voxels (64-CL-3x3), batch normalization (BN), rectified linear unit (ReLU), 128-CL-3x3, BN, ReLU, 256-CL-3x3, BN, ReLU, 1296 unit fully connected layer, BN, ReLU, spatial upsampling 2x2 (SU-2x2), 256-CL-3x3, BN, ReLU, SU-2x2, 128-CL-3x3, BN, ReLU, SU-2x2, 64-CL-3x3, BN, ReLU, 1-CL-1x1. A mean squared error loss function was used with the Adam optimizer to adjust network weights during training (Kingma and Ba, 2014) with a learning rate=0.0001. The 3D-DCA was trained for five epochs on the 119500 3D task-free fMRI volumes from the 100 subjects day 1 scans in our discovery dataset, with a batch size of 16 volumes and shuffling of the sequence of volumes in each epoch. This autoencoder was used to generate latent embeddings for all fMRI images in the study. For more details, see **Supplementary Methods**.

#### Latent space analysis

Latent space embeddings for all HCP scans were derived as the output from the 3D-DCA’s 1296-D middle layer. 216 dimensions had unvarying values across all volumes and were excluded from subsequent analysis, resulting in 1080 dimensions for each fMRI volume’s autoencoder middle layer code. The embeddings for the 119500 discovery dataset fMRI volumes were processed with Principal Component Analysis (PCA) in order to organize the latent space dimensions according to the amount of variance they explained. Matlab’s ‘pca’ function was run on the 119500 x 1080 embedding matrix with de-meaning of each column. The two-dimensional pairwise distributions for the first nine dimensions are shown in **Figure S3**. For gradient map validation, an independent latent space was derived for the 119500 validation dataset fMRI volumes. Partial Least Squares (PLS) regression was used to confirm that the same PCA components explaining maximal variance in the autoencoder embeddings also explained the most variance in the original BOLD timeseries. PLS regression was run on the 119500 x 1080 autoencoder embeddings against the 119500 x 273 cortical, subcortical, and cerebellar regional mean BOLD timeseries. To measure the similarity of the PCA and PLS components, the PCA-derived spatial gradient maps (see *Spatial gradient derivation*) were correlated with the PLS component spatial loadings.

#### Spatial gradient derivation

Following PCA, we regressed each gray matter voxel’s BOLD intensity values against timepoint-wise gradient slope (i.e. the PCA scores) to measure the strength of the linear relationship between latent space position and a given voxel’s BOLD activity. For a given component, the voxel activity/component beta values were derived for each voxel and stored in a statistical parametric “gradient” map (see **Supplementary Data 5**). We refer to these beta values as gradient weights. Independent gradient maps were derived for the validation dataset. These voxel-wise gradient maps were then averaged within each of a set of 273 regions of interest using a parcellation of 246 cortical and subcortical regions from the Brainnetome atlas (Fan et al., 2016) (http://www.brainnetome.org/) and 27 cerbellar regions from the SUIT atlas (Diedrichsen, 2006) (http://www.diedrichsenlab.org/imaging/suit.htm). Region names were assigned based on maximum probability overlap with the Harvard-Oxford cortical and subcortical atlas and the probabilistic cerebellar atlas in FSL. The polarity of each gradient was assessed by calculating what percentage of the 273 regions had positive or negative mean weights.

#### Timeseries analysis

Any voxel or region’s BOLD activity at a given timepoint can be reconstructed as the sum of the voxel/region’s gradient weights multiplied by the gradient slopes for the current timepoint, summed across all dimensions:

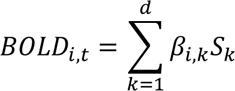

where *β*_*i,k*_ is the current voxel/region’s weight on a given gradient, *S* is the current timepoint’s gradient slope (ie PCA score) on a given dimension, *k* is the dimension number out of *d* total dimensions, *i* is the current voxel/region, and *t* is the current timepoint. The strength of the relationship in BOLD signal between two regions, the functional connectivity, is typically represented as the covariance or correlation of two voxels’ or regions’ timeseries. Covariance is defined as the product of the two regions’ BOLD activity values at each timepoint, averaged over timepoints. The BOLD covariance can be determined by multiplying the gradient slope matrix for each timepoint elementwise by the gradient weight matrix, averaging across all timepoints, and taking the sum of the resulting matrix. In general, the covariance between a pair of regions for a latent space with any number of dimensions is:

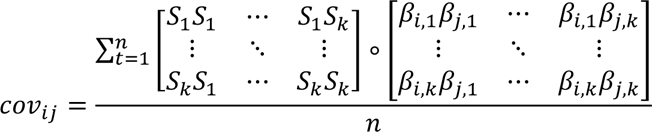

Where *Sk,t* is the gradient slope of fMRI timepoint *t* for latent dimensions 1 through *k*, and *β* is the gradient weight for region *i* or *j* on dimension *k*. In order to compute the correlation from the covariance, the standard deviation is required. The standard deviation for a given region is calculated by multiplying the gradient slope covariance matrix by the region *i*-region *i* gradient weight matrix:

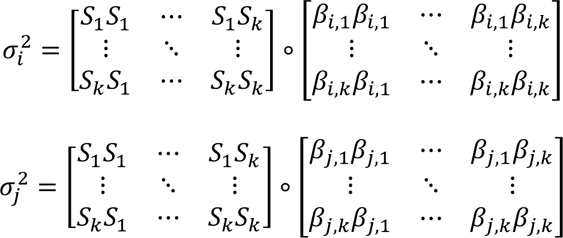

Finally, the correlation between region *i*-region *j* is determined by:

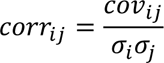

#### Activation fields

Because regional activation can be estimated using the region’s weight on each gradient and the slopes of those gradients, regional activation can be visualized for two dimensions of latent space as an “activation field” (**Figure 3A**). The overall BOLD activity level of a region at a specific point in a latent trajectory can be determined by its (*x*,*y*) position on the regional activation field. In order to visualize how a trajectory relates to functional connectivity (co-activation) between two areas, we applied the covariance formula. Covariance is measured as the mean of the product of two BOLD signals over time, assuming each signal has zero mean. The “instantaneous” covariance is thus the product of the BOLD values in two different regions in a given timepoint, which can be represented in latent space as the element-wise product of the two regional activation fields, creating a “co-variation field” (**Figure 3A, right)**. The covariance across a run can then be measured by taking the sum of the trajectory locations for each timepoint on the co-variation field, and dividing by the total number of timepoints. The correlation coefficient can subsequently be derived by dividing the covariance by the product of the standard deviations for each regional BOLD signal, where standard deviation in latent space is equivalent to the mean trajectory width along a given dimension.

#### Within-subject reliability assessment

We determined within-subject similarity of latent fMRI trajectories from day 1 and day 2 for the 100 subjects in the validation dataset using gradient slope covariance matrices, a measure of “fingerprinting” (Finn et al., 2015). We reconstructed these using between 1-50 latent dimensions, by deriving the gradient slope covariance matrix as described in the *Timeseries analysis* section. Our goal was to determine the minimum latent space dimensionality at which reliable individual differences in functional activity became apparent. We used trajectories for the 100 validation subjects, projected into the discovery-dataset defined latent space, to mitigate the possibility of certain dimensions/gradients being unique to specific individuals. The gradient slope covariance matrix for each day 1/day 2 scan was compared with every other gradient slope covariance matrix using the geodesic distance metric (Venkatesh et al., 2020). The matrix from the same subject on the opposite day was assigned a rank between 1-100, and a match was identified if the subject’s corresponding matrix was the best fit. For each number of dimensions, we determined the median rank of the corresponding matrix and the number of matches (**Figure S6**).

#### Task fMRI analysis

Preprocessed task fMRI data for the four tasks from the HCP were analyzed (working memory, motor, language, emotion) (https://db.humanconnectome.org/data/projects/HCP_1200). We used left-right phase encoded scans. Subsequent postprocessing steps for the task fMRI data, which had not undergone FIX-ICA denoising, included removing the first 5 scans and then denoising by regressing out the 6 motion parameters, temporal derivatives, squares, the white matter timeseries extracted using a mask of the highest probability cortical white matter according to the FSL tissue prior mask, and the CSF timeseries extracted using a mask in the central portion of the lateral ventricles. Scans were bandpass filtered in the 0.008-0.15Hz frequency range to match the task-free state data, voxel-wise normalized across time to a mean of zero and a standard deviation of one, and spatially downsampled to 48x48x48 images with 4 mm^3^ resolution. These images were passed through the task-free state-trained autoencoder from the discovery sample and the subsequent autoencoder-dimension-to-PCA-component loadings to obtain the latent space projections.

When determining task-specific activation patterns, the following regressors were used for each task: for working memory, the 2-back and 0-back conditions, merging blocks from the faces, places, tools and body parts stimulus blocks (Barch et al., 2013); for the motor task, the active condition combining the right hand, left hand, right foot, left foot, and tongue blocks, and the fixation condition; for language, the story and math conditions; and for emotion, the faces and shapes conditions. Task condition block regressors were convolved with a hemodynamic response function using the ‘spm_get_bf’ function in SPM12 (https://www.fil.ion.ucl.ac.uk/spm/software/spm12/). A general linear model was then fit for each subject and task, where the gradient slope timeseries for a given latent dimension was the response variable and the two HRF-convolved task regressors were the predictors. The parameter estimates for each condition were calculated and then contrasted between conditions using a two-sample t-test across subjects. The t-statistic of the between-condition gradient slope differences was obtained for each latent dimension (as shown for nine dimensions in **Figure 5B**). This t-statistic vector was multiplied by the corresponding set of voxel-wise gradient weight maps and summed to obtain the gradient-based task contrast map, with units of BOLD standard deviations. The mean gradient slopes for each task for each condition are shown in **Figure 5B**, along with the 95% confidence intervals. The conventional voxel-wise general linear model task activation maps were derived by taking the preprocessed BOLD images and regressing each voxel’s timeseries against the task regressors and performing a subsequent two-sample t-test on the voxel-wise parameter estimates for each condition across subjects. Spatial correlations (Pearson r-values) were computed between the task GLM maps and the gradient-based task contrast maps based on between 1 and 100 dimensions, considering 23714 voxels in a gray matter mask with 4 mm^3^ resolution that covered cortical, subcortical, and cerebellar regions.

#### Functional modules

The task-free functional connectivity matrix was derived for the discovery dataset by 1) reconstructing each region’s timeseries based on a specific subset of gradients (see *Timeseries analysis*) and 2) correlating each region’s resultant timeseries in a pairwise manner. The functional connectivity matrix was then r-to-Z transformed. Graph theory analyses were run using the Brain Connectivity Toolbox (BCT; https://sites.google.com/site/bctnet/). Networks were thresholded to keep the strongest 10% of edges. Modularity was determined using the Louvain algorithm (Blondel et al., 2008) as implemented in BCT, running 1000 iterations with the default parameters, choosing the partition that maximized the Q value. Connector and provincial hubs were determined by computing each region’s within-module degree Z-score and participation coefficient. Regions in the top tertile of within-module degree Z-score and the bottom tertile of participation coefficient were identified as provincial hubs, while regions in the top tertile of participation coefficient and the bottom tertile of within-module degree Z-score were identified as connector hubs. The remaining regions were labeled as non-hubs. Each region’s neighbors were identified as the top 10 most highly functionally correlated regions.

Gradient weights for different regions were statistically compared in two different ways.

When only considering the two-dimensional latent space based on gradients 2 and 3 (**Figure 4**), the angle between each region’s [2x1] gradient weight vector was computed. To compare the mean neighborhood angle for provincial hubs, connector hubs, and non-hubs, we used a Watson-Williams test using pycircstat (Berens, 2009). When considering higher dimensional latent spaces defined by three or more gradients, region gradient weight vectors were compared to each other using the cosine similarity. Cosine similarity values for hub and non-hub regions were then statistically compared using a non-parametric Mann-Whitney test.

#### Genetic spatial correlation

We compared each gradient map to Allen Human Brain spatial gene expression patterns using the ‘abagen’ package (https://github.com/rmarkello/abagen) (Arnatkevičiūtė et al., 2019; Hawrylycz et al., 2012). We used default options including for donors (all), tolerance (2 mm), collapsing across probes (diff_stability), and intensity-based filtering threshold (0.5). Expression data was available for 261 out 273 Brainnetome and SUIT regions from both hemispheres for 15655 genes. All non-cortical regions were eliminated because of substantial differences in subcortical expression values, which would hamper brain-wide spatial correlation estimates, leaving 202 regions. Data-driven filtering was used to remove regions with outlying expression values. Using K-means clustering we identified an outlying cluster with 6 regions, which were removed to give the final [15655 x 196] matrix of expression values. For each of the first nine gradients in the discovery or validation datasets, we calculated the spatial Pearson correlation between each 196-region gene expression vector and the 196-region gradient weight vector.

We defined the statistical significance threshold as the Bonferroni corrected p-value of p=.05 / 15655 genes / 9 gradients=3.55 x 10^-7^. Furthermore, gradient/gene expression spatial correlations were only reported as significant if they replicated Bonferroni significance for at least one gradient in both the discovery and validation datasets.

The resultant set of 3804 significant genes were submitted to a Gene Ontology Enrichment Analysis using GOrilla (http://cbl-gorilla.cs.technion.ac.il/) (Eden et al., 2009), using a background of all 15655 genes and corrected for multiple comparison using the default FDR q-value < 0.1. GOrilla recognized 14952 genes, of which 13922 were associated with a gene ontology term.

The principal component of spatial gene expression was derived by performing PCA on the [15655 x 196] regional expression matrix and obtaining the PCA scores for each of the 196 regions on the first component.

#### Differential equation modeling

For each task-free scan in the discovery dataset, the first and second derivatives of the 1195 timepoint gradient slope timeseries were calculated using the ‘gradient’ function in MATLAB. In this section we refer to the first derivative of gradient slope as ‘velocity’, and the second derivative as ‘acceleration’. For gradients 1-9, linear regression was used to estimate a given gradient’s acceleration as a linear function of the other gradients’ slopes, velocities, and an intercept. The parameter estimates (which we refer to as coupling parameters) for the 19 terms from this regression - the position and velocity terms for each gradient and an intercept - were then used to define a system of 9 differential equations:

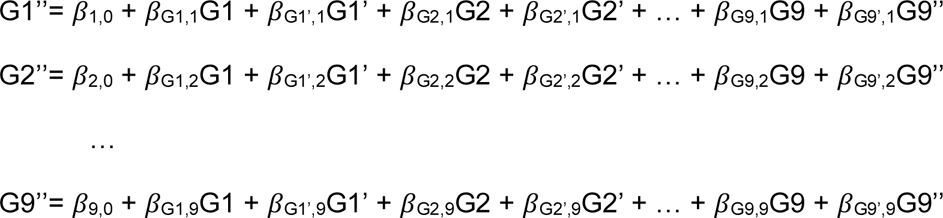

This system of equations was solved numerically using the ‘ode45’ function in MATLAB. For initial conditions, we used the gradient slopes/velocities for a given timepoint from the first 1170 timepoints from each scan. We determined each solution for an interval of 25 timepoints out from the initial condition (t=*0* to *t*=25) with a temporal resolution of 1 volume (720 ms). Forecasts were generated from each timepoint in the validation dataset, using coupling parameters derived from the discovery dataset. This yielded one 25x9 matrix for simulated gradient slopes, and a second 25x9 matrix for simulated gradient velocities. We compared the 25x9 simulated gradient slopes to corresponding true gradient slopes using correlation and root mean squared error to estimate the accuracy of the predictions.

For autoregressive modeling, we adapted the Matlab function from Liégeois and colleagues (Liégeois et al., 2017) to generate gradient timeseries forecasts (https://github.com/ThomasYeoLab/CBIG/blob/master/stable_projects/fMRI_dynamics/Liegeois2017_Surrogates/CBIG_RL2017_get_AR_surrogate.m). Parameters were derived in the discovery dataset task-free state data for each subject, and then averaged across subjects. These mean discovery parameters were then used generate gradient timeseries forecasts for each subject in the validation dataset, for a timeframe of 25 timepoints out from each actual timepoint in the task-free state scan. The variance explained r^2^ values were statistically compared between the differential equation and autoregression-based forecasts using the ‘corr_rtest’ function in MATLAB based on 117000 observations per group to derive Z-scores.

Simulations were performed using the system of differential equations for gradients 1-9. As an initial condition, we randomly selected 10000 timepoints from the task-free state scans in the discovery dataset. The associated initial conditions – each gradient’s slope, first derivative, and second derivative – were used as seeds to solve the differential equations. When solving the differential equations for simulations, the coupling parameters for each state – rest, working memory, motor, language, emotion – were estimated from the discovery dataset scans. Our basic strategy was to emulate a typical block-design task fMRI experiment. We did this by simulating 50-timepoint intervals from three different conditions: task-free, active task condition, or baseline task condition. Simulations were done for 50 timepoints with task-free coupling parameters. Next, simulations were done for a subsequent 50 timepoints with the coupling parameters from the active task condition (Working memory: 2-back; Motor: active; Language: story; Emotion: faces). A parallel simulation was run with 50 task-free timepoints followed by 50 timepoints with the coupling parameters from the baseline task condition (Working memory: 0-back; Motor: fixation; Language: math; Emotion: shapes). For each of the 10000 parallel active task or baseline task simulations, the last 50/100 timepoints were extracted and the first five of those timepoints were trimmed off to allow for stabilization. The remaining 45 timepoints were expanded from nine gradient timeseries into 273 region timeseries, from which 273x273 functional connectivity matrices were computed. These 10000 matrices were averaged to produce the active task condition-simulated functional connectivity (FC) matrix. For the actual data, the nine gradients’ timeseries from each timepoint where the subject was actively engaged in the task (ie where the HRF-convolved task waveform was greater than 0.5) were extracted from the validation dataset, expanded into 273 region timeseries, and pairwise correlated to obtain the actual active task condition FC matrix.

When comparing the average simulated and real matrices, we first statistically compared using ‘corr_rtest’ based on 37128 edges per matrix to derive Z-scores. To measure the specificity of the real and simulated matrices, we performed two additional analyses. First, we flattened the real and simulated FC matrices for five conditions – task-free state, working memory 2-back, motor active, language story, and emotion faces – into a 37128×10 r-to-Z transformed matrix and computed the 10×10 partial correlations. Second, we ran linear models predicting one of the 37128x1 columns using the other 37128x9 columns as predictors, then ran linear contrasts to determine if the parameter estimate for the real matrix for a given condition was significantly more similar to the corresponding simulated matrix than to any other matrix.

When forecasting task fMRI gradient timeseries using the differential equations, we first labeled volumes in each task as occurring during condition 1 when the condition 1 HRF-convolved task waveform was greater than 0.5. The same was done for condition 2, and the remaining volumes were labeled as baseline. The gradient slopes, velocities, and accelerations were then calculated separately for both conditions of each task for subjects in the discovery dataset. Gradient slope timeseries were forecasted the same way as for the task-free state data, limiting the prediction interval to 10 timepoints, after which prediction errors generally reached a plateau. For the main condition of interest in each task, three sets of forecasts were generated for each subject in the validation dataset: using coupling parameters derived from the same task condition, from the opposite task condition, or from task-free state data. The forecast accuracy was measured as the root mean squared error (RMSE) of actual and forecasted data at each timestep for each gradient, and then averaged across gradients at a given timestep.

The three sets of forecasts were statistically compared for accuracy using pairwise two-sample t-tests of mean RMSE at each timestep, corrected for multiple comparisons using the false discovery rate with q=0.05.

#### Data availability

Original data was obtained from the Human Connectome Project (1U54MH091657, PIs Van Essen and Ugurbil) and the Allen Human Brain Atlas (http://human.brain-map.org/). All code and processed data will be available upon publication at https://github.com/jbrown81/gradients

## Acknowledgements

This work was supported by NIH grant K01-AG055698 to JAB, K99-AG065457 to LP, and the Tau Consortium.

## Notes

### Competing Interest Statement

The authors have declared no competing interest.

