## Supplemental Information for "A dynamic gradient architecture generates brain activity states"

**Appendix**

Supplementary Results

Supplementary Methods

Supplementary References

Supplementary Tables 1-4

Supplementary Figures 1-10

Supplementary Movie 1

**Supplementary Results**

*Low-dimensional latent space explains majority of brain activity variance and is reproducible*

We first examined the intrinsic dimensionality of the task-free state fMRI-derived latent space. PCA was performed on the 119500 x 1080 autoencoder codes. The first five PCA components explained 22.5% of variance in the discovery dataset and 142 components explained 66% of the variance. For comparison to PCA, Partial Least Squares (PLS) regression was used to find components maximizing the co-variation between the 119500 x 1080 autoencoder codes and the 119500 x 273 cortical, subcortical, and cerebellar regional mean BOLD timeseries. The first three PLS components explained 44.9% of BOLD variance across regions (44%±16% in cortical areas, 15%±7% in subcortical areas, 28%±13% in cerebellar areas) **(Figure S1)**. Overall, gradient maps explained the most variance in cortical areas, intermediate levels in cerebellar areas, and the least variance in subcortical areas (**Figure S2)**. Five components were needed to explain more than half of the variance (51%) and all 1080 components explained 90% of the variance. The PCA components corresponded well with the PLS-derived components (PCA component 1 variance explained=10.2%; correlation with PLS component 1 r=0.999; PCA component 2: variance=4.1%, correlation with PLS component 2 r=0.995; PCA component 3: variance=3.6%, correlation with PLS component 3 r=0.996; see full results in **Table S4**). With both PCA and PLS, the first component explained substantially more variance than any next highest component (10.2%/2.5x and 32.5%/4.36x, respectively). Importantly, the 4th PCA component in the discovery dataset did not have a top match with any of the PLS components. Based on this information and further investigation (see Gradient reproducibility), this component was excluded from subsequent analysis and dimensions 5 and higher were promoted by one rank. All subsequent analysis was based on the PCA components. We focused most analyses on the first nine components based on the criterion that this was where the second derivative of the explained variance curves became positive, indicating a diminishing return of subsequent variance explained.

*Gradient reproducibility*

Spatial gradient maps were highly reproducible in the validation dataset. For the first three components from the discovery dataset, the spatial correlation of the spatial gradients with the corresponding component gradients from the validation set were high (r=0.96, r=0.84, r=.80). The first 12 dimensions appeared in nearly the same sequence with a few minor variations **(Table S1 and Figure S4/Figure S5)** and spatial correlation coefficients above r=0.5. After the 12th discovery gradient, the spatial correlations fell below r=0.5. The most noteworthy difference between the discovery and validation datasets was the failure to replicate discovery component 4 and its associated gradient map in the validation dataset. Upon further examination, the PCA component underlying gradient 4 had very small number of autoencoder dimensions loading on it. We calculated the skewness of the autoencoder loading values for each component and found that discovery component 4 had a skewness 4.45 times more extreme than any other component in the dataset, while in the validation dataset no component had skewness more than 1.80x. A subject-focused analysis found that one discovery subject (subject 66) loaded much more heavily onto component 4 and the underlying autoencoder dimensions than any other subject (component mean loading Z=7.06; no other subject loading Z > 1.85). Thus while this subject could be considered an outlier, we chose to leave them in the analysis due to their inclusion in the HCP dataset. Instead, we opted to exclude discovery component 4 from subsequent analysis based on 1) its heavy skew towards one outlying subject, 2) its lack of a best-fit match with any PLS component, and 3) its lack of replication in the validation dataset. The other noteworthy discovery/validation gradient difference was the splitting of discovery dataset gradient 2 into two similar gradients (2 and 3) in the validation dataset. Such dimension splitting may be driven by individual-specific anatomical variation (ie phenotypic differences) or differences in functional properties of the gradient (ie state differences).

*Latent space trajectories reflect shifting gradient slopes*

The majority of fMRI volumes appeared near the origin of latent space, with a progressively lower percentage of volumes occurring farther from the origin **(Figure S3)**. This was expected because each fMRI run was demeaned across time for each voxel before training the autoencoder. When qualitatively examining the relative positions of all volumes within a given subject's fMRI scan, adjacent volumes always appeared in close proximity **(Figure 1A and Movie S1)**. This latent space proximity and continuity occurred despite the autoencoder not having any information about subject identity or the temporal relationship between volumes. When volumes were assembled into latent space trajectories, different individuals exhibited substantial variability in the width and mean direction of transitions (see **Figure 3**).

*Co-variation between regions is determined by pairwise interactions of gradients*

Our next goal was to understand how interactions between gradients, as captured in latent space trajectories, relate to different levels of regional activation and inter-regional co-variation. Because activation in a region can be estimated using the region's weight on each gradient and the slope of that gradient, regional activation can be represented in two dimensions as an activation "field" in latent space **(Figure 3A)**. We assessed activation and co-variation in the anterior cingulate cortex (ACC) and the middle frontal gyrus (MFG). The ACC had a near-zero weight on gradient 2 and a large negative weight on gradient 3, resulting in an activation field that was most positive for more negative values on dimension 3, corresponding to negative gradient 3 slopes. MFG had a positive weight on gradient 2 and a negative weight on gradient 3, and consequently had an activation field with the most positive BOLD values where there was a combination of highly positive values on dimension 2 (positive gradient 2 slope) and highly negative values on dimension 3 (negative gradient 3 slope). The co-variation field, the product of the two activation fields, had maxima in the upper left and lower right quadrants of latent space, with an optimal angle at the midpoint between the trajectory directions to maximize activity in either the ACC or MFG. In order to maximize covariation, a trajectory simply has to maximally access these high co-variation areas of latent space. In order to maximize BOLD correlation, two criteria must be balanced: the trajectory should maximize co-variation (the numerator of the correlation formula) while having a narrow trajectory width orthogonal to the optimal co-variation angle (minimizing the standard deviations, the denominator of the correlation formula). As stated in the main results (**Figure 3B**), the individual subjects with maximal or minimal ACC-MFG correlation had trajectories that were most or least extended along the optimal co-variation angle for this pair of regions, and had relatively narrow trajectories running either along the co-activation global maximum or perpendicular to it. The subject trajectories with the most positive and most negative ACC-MFG covariation rather than correlation are shown in **Figure S7**. This illustrates that different types of trajectories are required to maximize the covariance versus the correlation, in line with previous work suggesting correlation is not necessarily preferable to covariance as a measure for detecting differences between individuals (1).

*Within-subject reliability assessment*

For the 100 subjects in the validation dataset, the median rank of the match rank between the day 1 and day 2 gradient slope covariance matrices was one with six dimensions (**Figure S6)**.

The number of matches reached 100% with 29 dimensions. Thus, a small number dimensions were sufficient to represent an identifiable fingerprint of brain activity in most subjects, while a longer tail of dimensions were necessary to reach perfect sensitivity and specificity.

*Basis for functional modularity and hubness*

When considering functional connectivity based on the first 10 gradients, seven modules were detected and provincial hubs had significantly smaller cosine distance to their most strongly connected neighbors than non-hubs (provincial hubs: mean=0.11±0.07; non-hubs: mean=0.14±0.09; Mann-Whitney U=349598.5, $p=3.19x10^{-08}$; **Figure S8**), which in turn had smaller cosine distance to their neighbors than connector hubs (connector hubs: mean=0.19±0.09; Mann-Whitney U=198090.0, $p=6.46x10^{-25}$). With 100 gradients, six modules were detected and provincial hubs still had significantly smaller cosine distance to their neighbors (provincial hubs: mean=0.32±0.15; non-hubs: mean=0.41±0.17; Mann-Whitney U=281320.5, $p=1.27x10^{-27}$; **Figure S8**), and non-hubs to had significantly smaller cosine distance than connector hubs (connector hubs: mean=0.48±0.15; Mann-Whitney U=179698.0, $p=6.04x10^{-09}$).

*Underlying gene expression patterns*

For each gradient, the number of genes with significant spatial correlations was: gradient 1: 3572, gradient 2: 48, gradient 3: 0, gradient 4: 3, gradient 5: 387, gradient 6: 15, gradient 7: 50, gradient 8: 0, gradient 9: 14.

*Predicting dynamic trajectories during tasks*

When reconstructing based on the state-specific coupling parameters during the working memory task, forecasts were accurate for around 20 timepoints before becoming unstable. Importantly, these reconstructions were more accurate when including all nine gradients in a system of equations with terms for mutual influence rather than when modeling each gradient independently. The reconstruction of the group-averaged gradient timeseries for the working memory 2-back condition was compared when using a system of differential equations for gradients 1-9 versus when modeling each gradient with an independent equation, where the only parameters were the gradient's own slope and first derivative. The system of equations was more accurate, with a difference in variance explained by reconstructions (system of equations versus independent equations) for timepoints 1-30 of $r^2$=0.00, 0.01, 0.02, 0.02, 0.03, 0.03, 0.03, 0.04, 0.05, 0.07, 0.10, 0.13, 0.13, 0.08, 0.10, 0.30, 0.56, 0.72, 0.78, 0.75, 0.67, 0.60, 0.53, 0.47, 0.38, 0.27, 0.18, 0.28, 0.12, -0.05. Thus, the system of equations outperformed the independent equations particularly on the on timepoints farther out from the initial condition, here timepoints 16-25. In general, we observed a tradeoff where coupling parameters estimated from a small number of timepoints (<100) could generate accurate reconstructed timeseries for a short timeframe before exponentially exploding. Conversely, parameters estimated from larger samples made slightly less accurate short-run forecasts but with greater long-run stability.

The simulated timeseries for the active condition of each task were generated with state-matched coupling parameters from the discovery dataset. In the specificity analysis, we found that the partial correlations and F-statistics for the simulated FC matrix from rest and each task were always most similar to the corresponding real matrix (rest: partial r=0.96, F=180740; working memory: partial r=0.70, F=1220; motor, partial r=0.63, F=5010; language, partial r=0.62, F=425; emotion, partial r=0.61, F=2390; all $p < 0.00001$). This suggests that the simulated FC dynamics are sufficiently realistic to yield FC matrices that are indistinguishable from actual FC matrices, and that even with variable initial conditions the simulated trajectories converge on the same mode of gradient interaction.

In each of the four tasks, the coupling parameters from the condition of interest derived from the discovery dataset yielded more accurate forecasts in the validation dataset than when using parameters from the opposite condition or from task-free state parameters. For working memory, 2-back task-derived parameters were more accurate than 0-back-derived and rest-derived parameters for times $t$=1-10 (all FDR-corrected $p < 0.01$, all $t > 3$). During the motor task, parameters derived from the union of all motor conditions were more accurate than fixation-derived parameters for times 5-10 and rest-derived parameters for times $t$=1-10 (all $p < 0.01$, all $t > 3$). For language, parameters from the 'story' condition were more accurate than math condition-derived and rest-derived parameters for times $t$=1-10 (all $p < 0.01$, all $t > 3$). In the emotion task, coupling parameters from the 'faces' condition were more accurate than shapes condition-derived and rest-derived parameters for times $t$=1-10 (all $p < 0.01$, all $t > 3$).

### Supplementary Methods

*Subjects and data*

All MRI data used in this study were publicly available and anonymized. Informed consent was obtained for each individual by the HCP consortium and were previously approved by the Washington University Institutional Review Board as part of the HCP. The HCP complied with all relevant ethical regulations. The present study agreed to the Open Access Data Use Terms (https://www.humanconnectome.org/study/hcp-young-adult/document/wu-minn-hcp-consortium-open-access-data-use-terms) and was exempt from the UCSF IRB because investigators could not readily ascertain the identities of the individuals to whom the data belonged. Task-free state scans were 14.4 minutes long with a repetition time (TR) of 720 ms, resulting in 1200 fMRI volumes per scan. Only the left-right phase encoded scans were used. FSL (https://fsl.fmrib.ox.ac.uk/fsl/fslwiki/) and AFNI (https://afni.nimh.nih.gov/) were used for subsequent fMRI preprocessing. The first 5 volumes for each fMRI scan were dropped to allow scanner stabilization. Scans were bandpass filtered in the 0.008-0.15Hz frequency range and subsequently normalized in each voxel across time to have zero mean and unit variance.

*Deep learning*

A 3D-DCA is a deep neural network with two parts, an encoder and a decoder. It is designed to learn a compressed representation of an input image through progressively smaller sets of features (the encoder), then progressively decompress features (the decoder) into a reconstruction of the original image that should match the input as precisely as possible. The middle layer of the autoencoder is the information “bottleneck” and the output activations from this layer are latent space embeddings. The primary advantage 3D-DCAs offer over other dimensionality reductions methods such as PCA, ICA, and tSNE is that when applied to 3D MRI data, an autoencoder is explicitly designed to learn spatial 3D features. The 3D-DCA considers spatially adjacent 3x3 patches of voxels in lower layers of the encoder, and then combines these lower-order features into spatially distributed features at the highest layer of the encoder.

Autoencoder setup and training was performed using Keras (https://keras.io/) with a Tensorflow backend (https://www.tensorflow.org/) on an Amazon AWS P2 instance with a NVIDIA K80 GPU. After 5 epochs and 36 hours of model training, the post-epoch mean squared error in the training sample decreased from 0.1606 standardized BOLD intensity units after the first epoch to 0.1422 after the fifth epoch and reached a plateau. The error in the validation dataset correspondingly decreased from 0.1692 to 0.1473, indicating that the 3D-DCA was able to accurately reconstruct unseen images outside of the training set. We used this autoencoder to generate latent embeddings for all fMRI images from the HCP dataset. For the purpose of gradient validation, a second 3D-DCA was trained independently on the 119500 task-free fMRI volumes from the 100 subjects in the validation dataset.

*Timeseries analysis*

In a latent space representation, shown in an example here for two hypothetical regions on two dimensions, if the BOLD signal formulas for the two regions are the following:

Region 1: BOLD signal(time $t$) = 3$a$ + 2$b$

Region 2: BOLD signal(time $t$) = 1$a$ + 4$b$

Where $a$ and $b$ are the fMRI volume gradient slopes at time t, and the constants region gradient weights.

These can be multiplied by each other to obtain the BOLD covariation. The multiplication of the two formulas can expanded using the distributive property:

$(3a + 2b) * (1a + 4b) = 3a^2 + 12ab + 2ba + 8b^2$

This co-activation formula can be represented as a matrix:

$[3a^2\ 12ab$

$2ba\ 8b^2]$

The sum of these matrix elements, when calculated for this timepoint with the current gradient slopes for $a$ and $b$, is the (instantaneous) region $a$-region $b$ covariance. These matrices can be averaged across all timepoints and summed to determine the overall covariance. This matrix can be decomposed into two elements, 1) the gradient slope covariance matrix (i.e. the outer product),

*Functional modules*

The BOLD timeseries for each region was reconstructed for the 273 regions. Latent projections of task-free fMRI data were concatenated for all subjects in the discovery dataset. For the modularity analysis, functional connectivity matrices were derived based on a subset of gradients by 1) multiplying each region's gradient weight by the gradient slope for each timepoint and summing across the set of included dimensions (ie the dot product) and 2) correlating each region's resultant timeseries in a pairwise manner. The functional connectivity matrix was then r-to-Z transformed. Graph theory analyses were run using the Brain Connectivity Toolbox (BCT; https://sites.google.com/site/bctnet/). Networks were thresholded to keep the strongest 10% of edges. Modularity was determined using the Louvain algorithm (4) as implemented in BCT, running 1000 iterations with the default parameters, choosing the partition that maximized the Q value. Connector and provincial hubs were determined by computing each region's within-module degree Z-score and participation coefficient. Regions in the top tertile of within-module degree Z-score and the bottom tertile of participation coefficient were identified as provincial hubs, while regions in the top tertile of participation coefficient and the bottom tertile of within-module degree Z-score were identified as connector hubs. The remaining regions were labeled as non-hubs. Each region's neighbors were identified as the top 10 most highly functionally correlated regions.

$$G1'' = \beta_{1,0} + \beta_{G1,1}G1 + \beta_{G1',1}G1' + \beta_{G2,1}G2 + \beta_{G2',1}G2' + \ldots + \beta_{G9,1}G9 + \beta_{G9',1}G9''$$

$$G2'' = \beta_{2,0} + \beta_{G1,2}G1 + \beta_{G1',2}G1' + \beta_{G2,2}G2 + \beta_{G2',2}G2' + \ldots + \beta_{G9,2}G9 + \beta_{G9',2}G9''$$

$$\ldots$$

$$G9'' = \beta_{9,0} + \beta_{G1,9}G1 + \beta_{G1',9}G1' + \beta_{G2,9}G2 + \beta_{G2',9}G2' + \ldots + \beta_{G9,9}G9 + \beta_{G9',9}G9''$$

**Supplementary Tables**

| Discovery set dimension gradient | Best matched validation set dimension gradient | Spatial correlation (Spearman r) |
|---|---|---|
| 1 | 1 | 0.96 |
| 2 | 2/3 | 0.66/0.84 |
| 3 | 4 | .80 |
| 4 | 5 | 0.74 |
| 5 | 6 | 0.79 |
| 6 | 7 | 0.60 |
| 7 | 8 | 0.74 |
| 8 | 9 | 0.62 |
| 9 | 12 | 0.58 |
| 10 | 11 | 0.59 |
| 11 | 10 | 0.55 |

**Table S1.** Spatial correspondence between latent dimension spatial gradients derived from the discovery and validation datasets where r > 0.5.

| Process | Function | Component |
|---|---|---|
| metal ion transport | metal ion transmembrane transporter activity | transmembrane transporter complex |
| inorganic cation transmembrane transport | ion gated channel activity | transporter complex |
| inorganic ion transmembrane transport | gated channel activity | ion channel complex |
| ion transmembrane transport | inorganic molecular entity transmembrane transporter activity | cation channel complex |
| ion transport | ion transmembrane transporter activity | plasma membrane |

| | | |
|---|---|---|
| cation transmembrane transport | transporter activity | plasma membrane protein complex |
| regulation of hormone levels | inorganic cation transmembrane transporter activity | plasma membrane part |
| regulation of transport | calcium ion binding | intrinsic component of plasma membrane |
| regulation of localization | transmembrane transporter activity | cell projection |
| potassium ion transport | cation channel activity | voltage-gated potassium channel complex |
| cation transport | ion channel activity | integral component of plasma membrane |
| potassium ion transmembrane transport | delayed rectifier potassium channel activity | plasma membrane bounded cell projection |
| cellular potassium ion transport | substrate-specific channel activity | potassium channel complex |
| signaling | passive transmembrane transporter activity | neuron part |
| regulation of system process | channel activity | neuron projection |
| export across plasma membrane | voltage-gated channel activity | extracellular space |
| system process | voltage-gated ion channel activity | membrane part |
| regulation of ion transport | cation transmembrane transporter activity | synapse |
| regulation of transmembrane transport | voltage-gated cation channel activity | intrinsic component of membrane |
| transmembrane transport | potassium ion transmembrane transporter activity | integral component of membrane |
| cell communication | voltage-gated potassium channel activity | extracellular matrix |
| anatomical structure development | ATPase activity, coupled to transmembrane movement of ions, phosphorylative mechanism | voltage-gated sodium channel complex |

**Table S2.** The top 22 ranked terms from the gene ontology enrichment analysis, based on Allen Human Brain Atlas gene expression map spatial correlation with gradient maps 1-9.

| Simulated FC matrix | WM 2-back | WM 0-back | Motor active | Motor fixation | Language story | Language math | Emotion faces | Emotion shapes |
|---|---|---|---|---|---|---|---|---|

| | | | | | | | | |
|---|---|---|---|---|---|---|---|---|
| Working memory 2-back | **0.95** (Z= 12.09) | 0.94 | 0.85 | 0.92 | 0.87 | 0.91 | 0.92 | 0.90 |
| Motor active | 0.80 | 0.81 | **0.94** (Z= 5.33) | 0.92 | 0.80 | 0.87 | 0.91 | 0.93 |
| Language story | 0.91 | 0.92 | 0.87 | 0.91 | **0.93** (Z=7.83) | 0.91 | 0.91 | 0.90 |
| Emotion faces | 0.87 | 0.87 | 0.87 | 0.94 | 0.84 | 0.89 | **0.95** (Z=21.45) | 0.94 |

**Table S3.** Correspondence between simulated data FC matrices for each active task condition and the actual task FC matrices. All correlations are r values and for the strongest correlation in each row, the Z-statistic from statistical comparison of the highest and second highest correlation coefficients is shown (all $p < 0.001$ based on 37128 edges per FC matrix).

| PLS component | Best matched PCA component | Spatial correlation |
|---|---|---|
| 1 | 1 | 0.999 |
| 2 | 2 | 0.995 |
| 3 | 3 | 0.996 |
| 4 | 6 | 0.901 |
| 5 | 5 | 0.804 |
| 6 | 8 | 0.913 |
| 7 | 7 | 0.864 |
| 8 | 6 | 0.497 |
| 9 | 9 | 0.817 |
| 10 | 10 | 0.791 |
| 11 | 11 | 0.849 |
| 12 | 12 | 0.661 |
| 13 | 15 | 0.618 |
| 14 | 14 | 0.421 |
| 15 | 13 | 0.612 |
| 16 | 18 | 0.494 |
| 17 | 17 | 0.399 |
| 18 | 20 | 0.315 |
| 19 | 2 | 0.275 |
| 20 | 19 | 0.438 |

**Table S4.** Spatial correlation between the PCA-derived gradient maps and the PLS-derived spatial loadings maximizing covariance between the autoencoder codes and the 273 regional BOLD timeseries.

**Supplementary Figures**

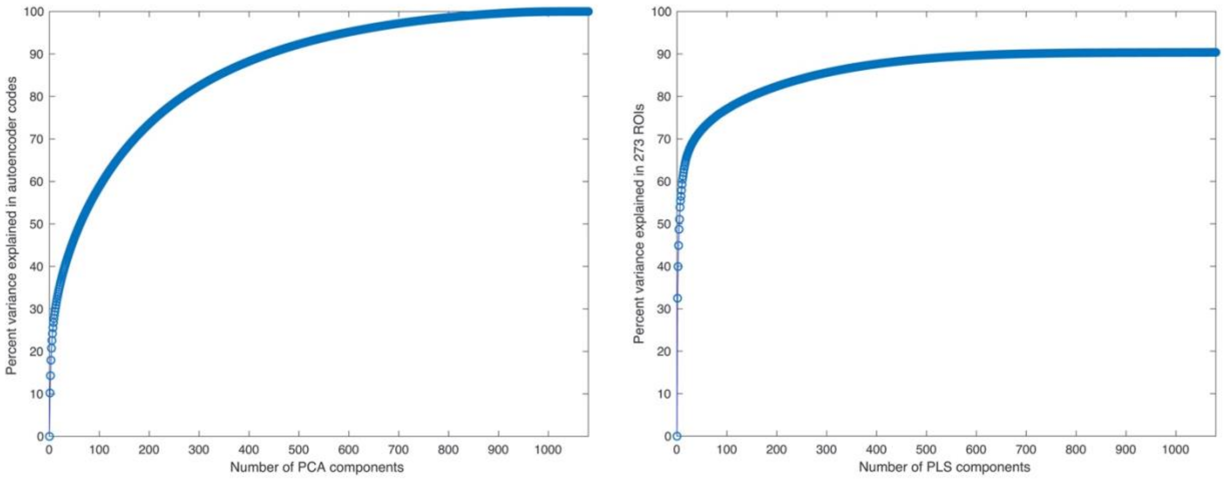


**Figure S1. Percent of variance explained by latent space components.** Left: the amount of cumulative variance explained by the first 100 PCA components derived from the 119500 x 1080 autoencoder codes. Right: the amount of cumulative variance explained by the first 100 PLS components relating the autoencoder codes to the 119500 x 273 region BOLD timeseries.

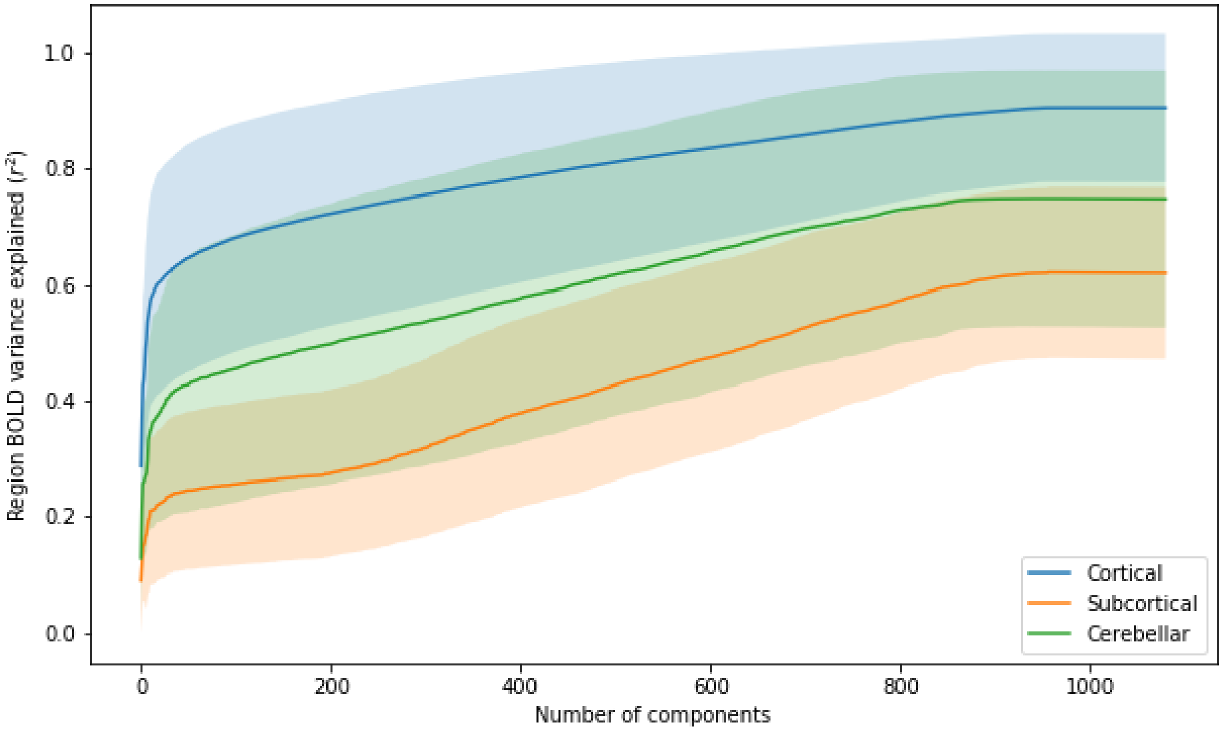


**Figure S2. Region BOLD signal variance explained by latent dimensions.** Lines show mean variance explained by the PCA-defined latent dimensions for the 273 regions that were cortical (blue), subcortical (orange), or cerebellar (green). Shaded bands show the standard deviation.

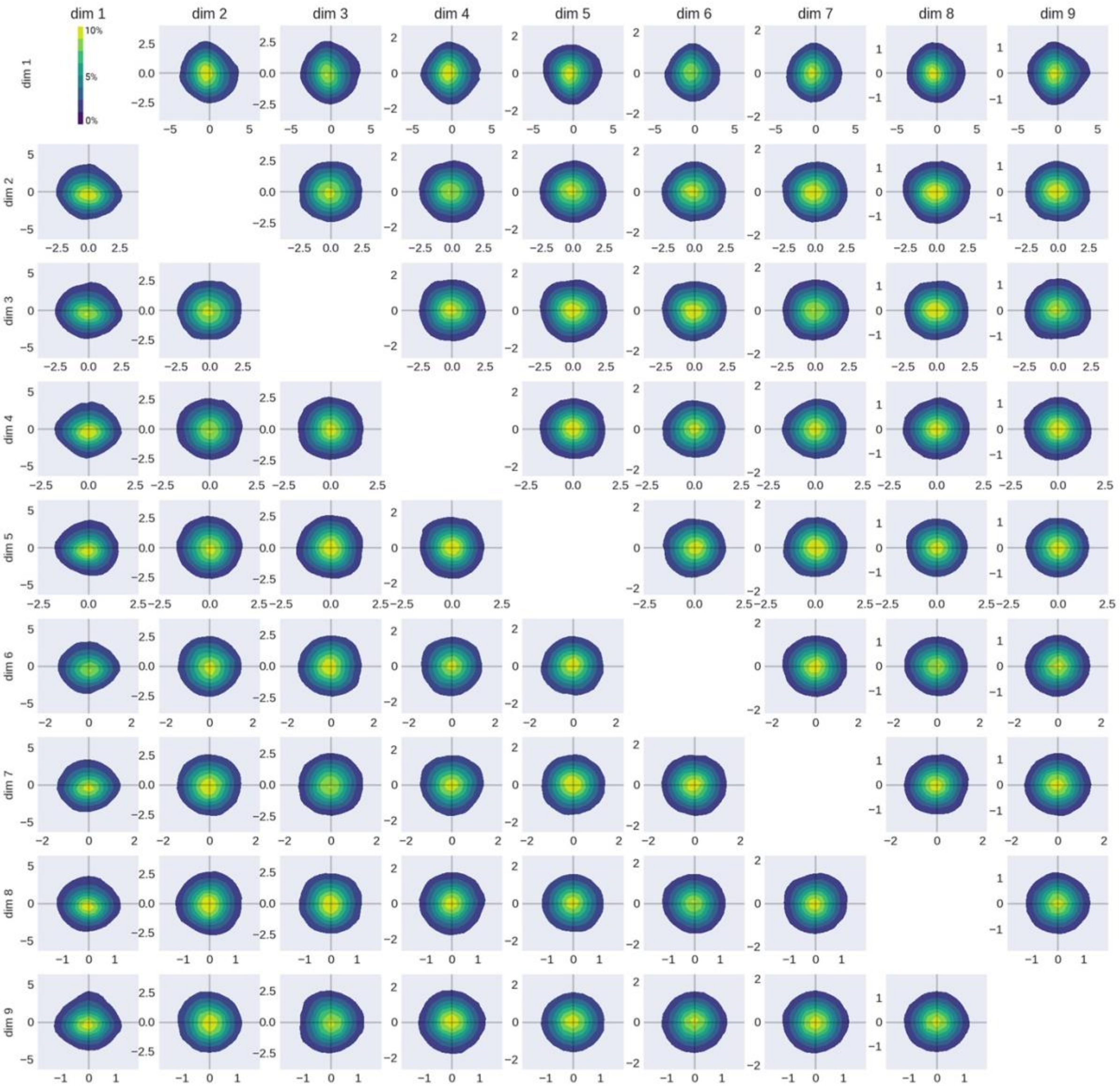


**Figure S3. Latent space two-dimensional density plots.** The pairwise dimension density plots show the percentage of fMRI volumes from the discovery dataset (n=119500) that occur in different latent space positions. Dimensions limits are based on ±3 standard deviations for each dimension.

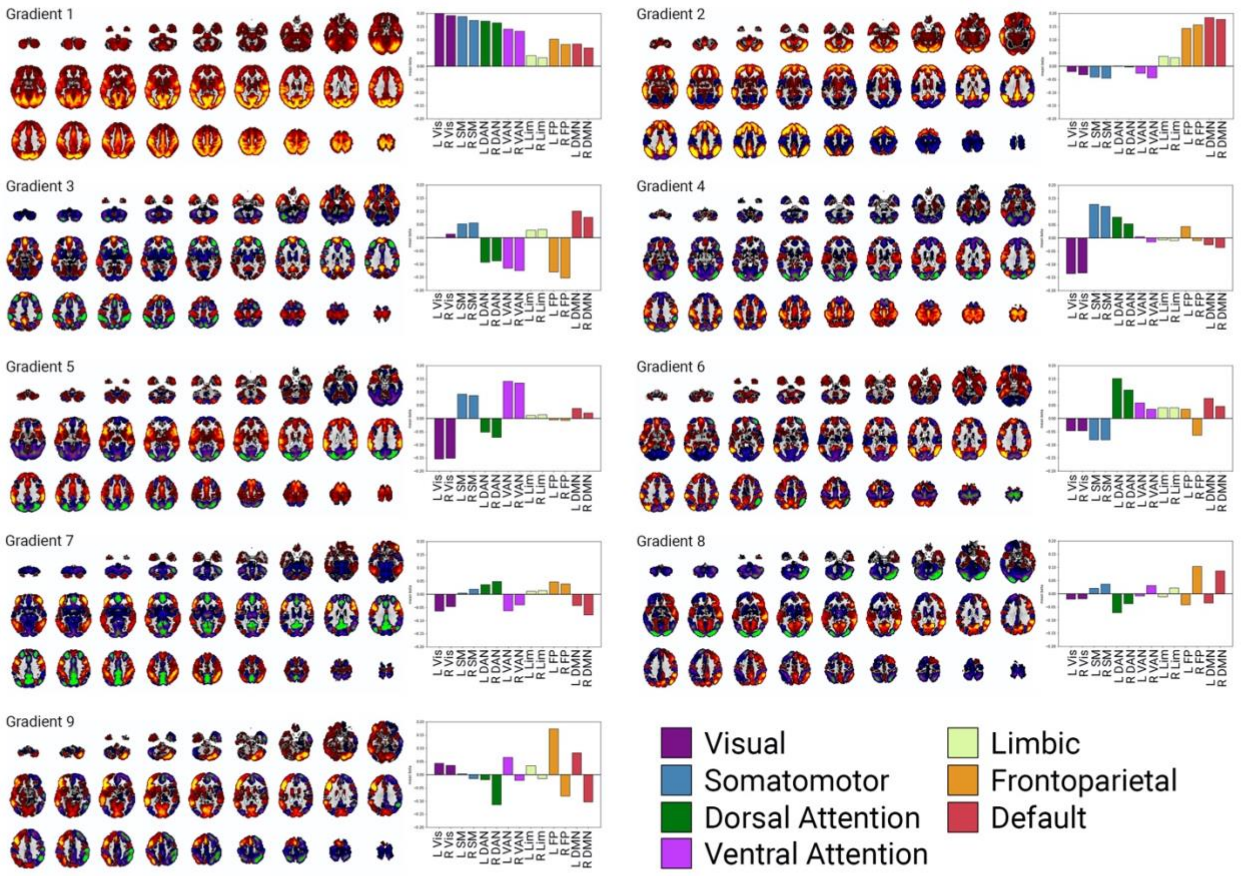


**Figure S4. Discovery dataset gradient map correspondence with functional connectivity networks.** Gradient maps for dimensions 1-9 in the discovery dataset. Bar plots show the mean beta weight for voxels in each gradient map overlapping with each of 7 cortical networks in the left or right hemisphere.

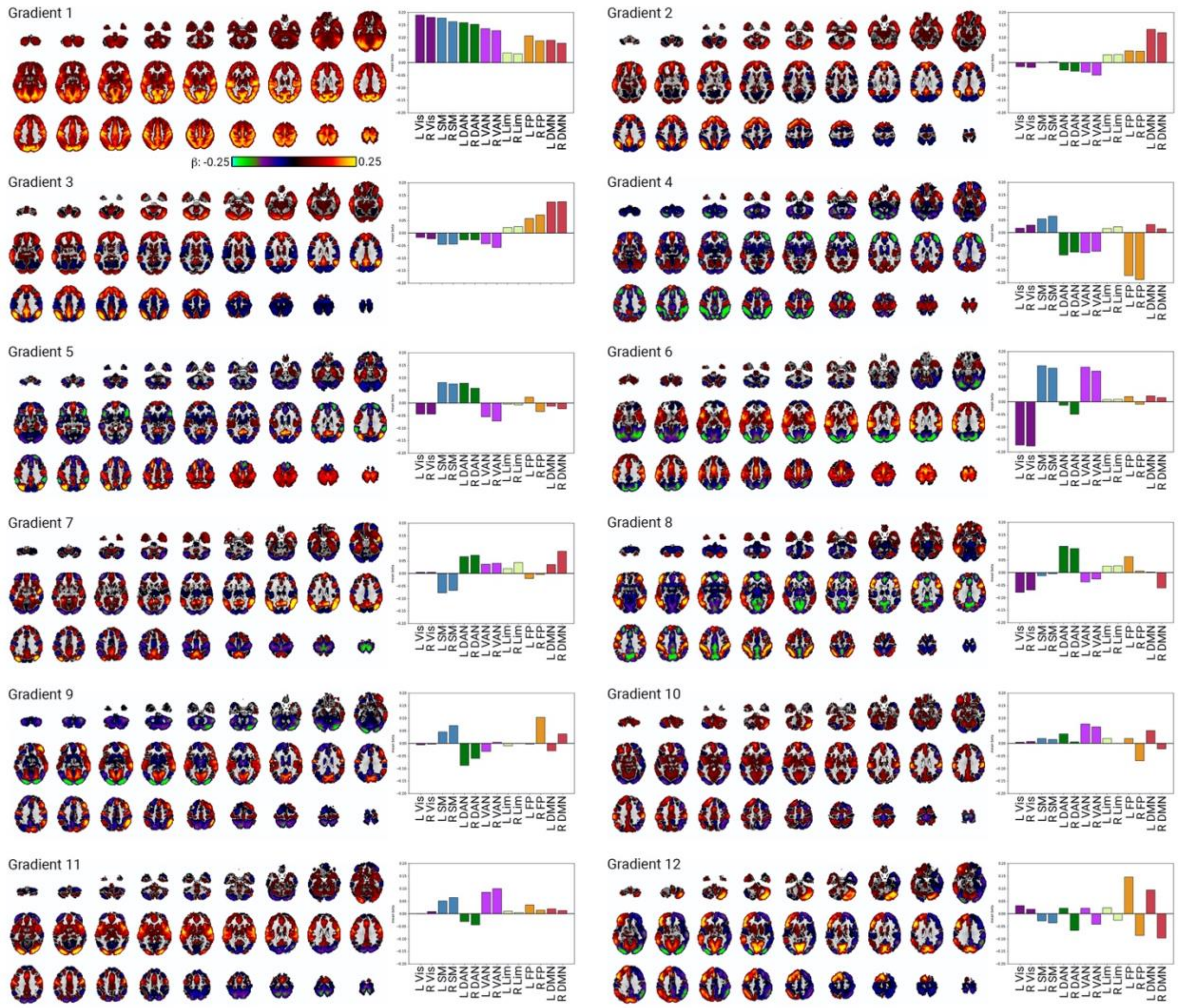


**Figure S5. Validation dataset gradient map correspondence with functional connectivity networks.** Gradient maps for dimensions 1-12 in the validation dataset, showing the best match for each discovery gradient (see **Table S1**). Bar plots show the mean beta weight for voxels in each gradient map overlapping with each of 7 cortical networks in the left or right hemisphere.

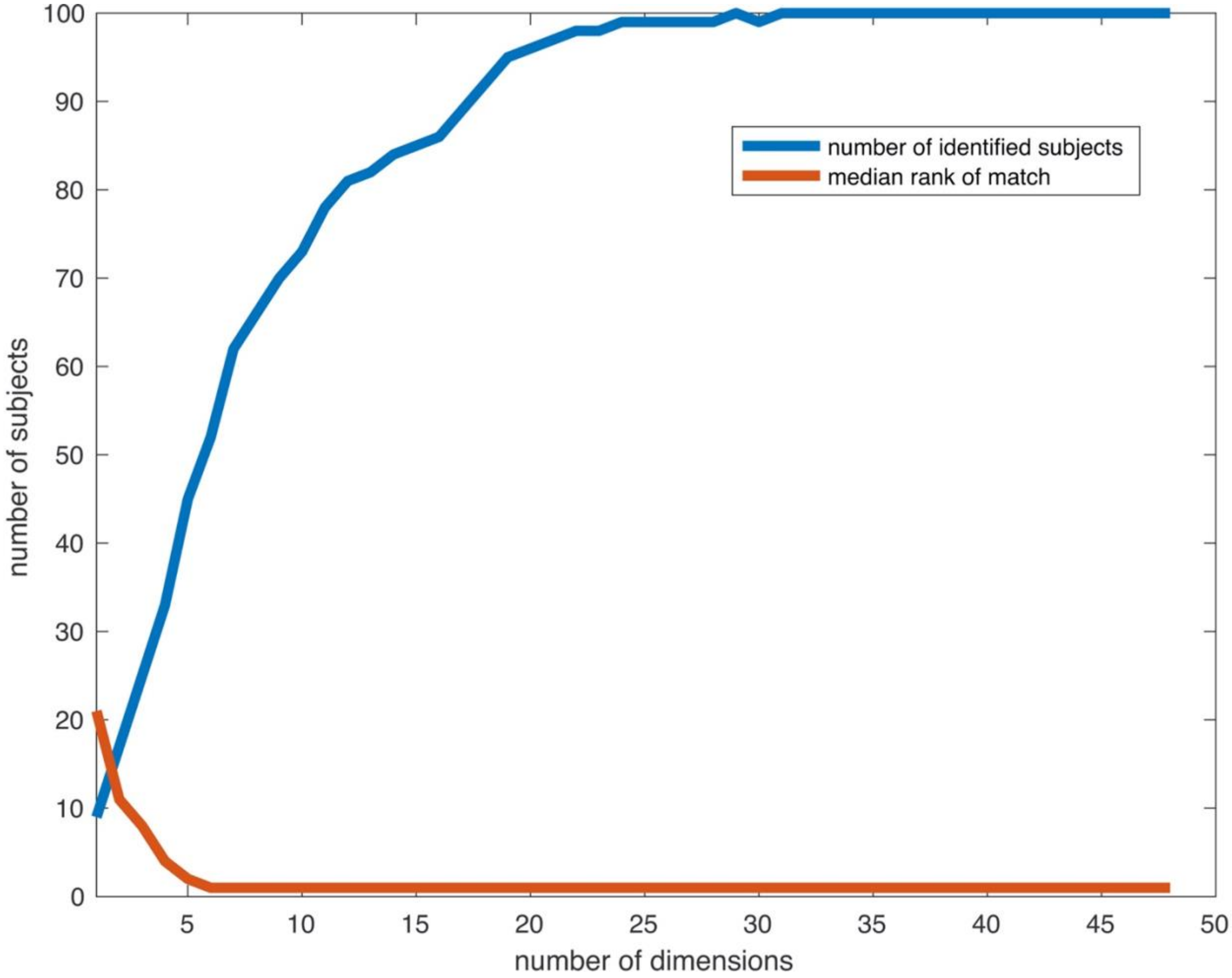


**Figure S6. Reliability of latent trajectories.** Within-subject reliability of gradient slope covariance matrices, from day 1 and day 2 comparisons for 100 subjects from the validation dataset, based on the number of latent space dimensions. Plots show the number of subjects for whom their own day 2 scan was the top match for their day 1 scan (blue) and the median similarity rank of a subject's day 2 scan to their day 1 scan (out of 100; orange).

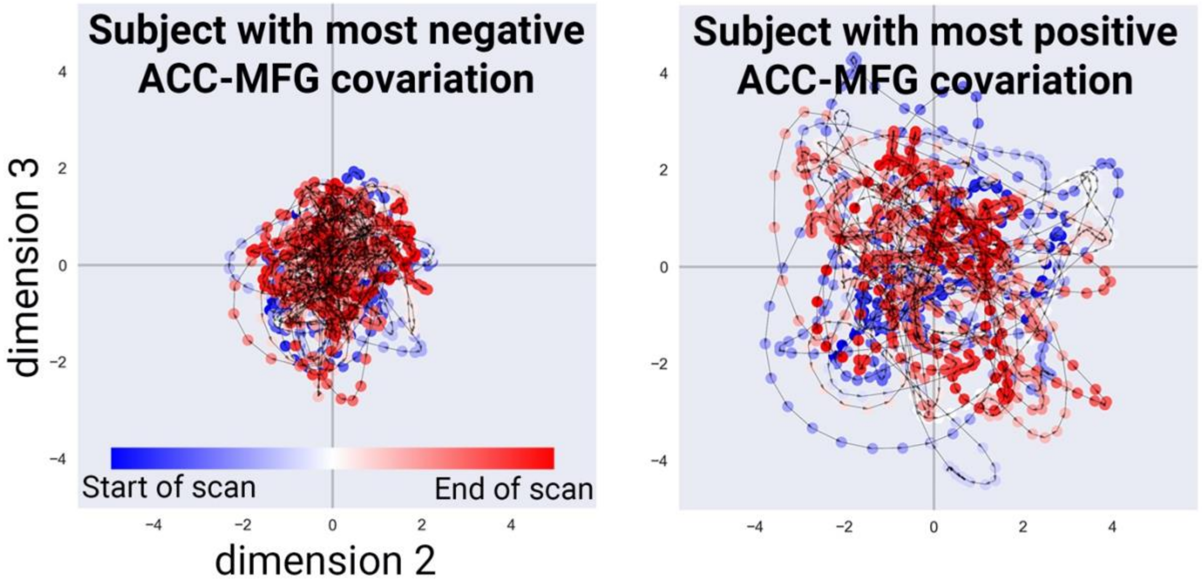


**Figure S7. Trajectories maximizing covariance between specific regions.** Latent trajectories during task-free state for subjects that have the most negative or most positive ACC-MFG covariation.

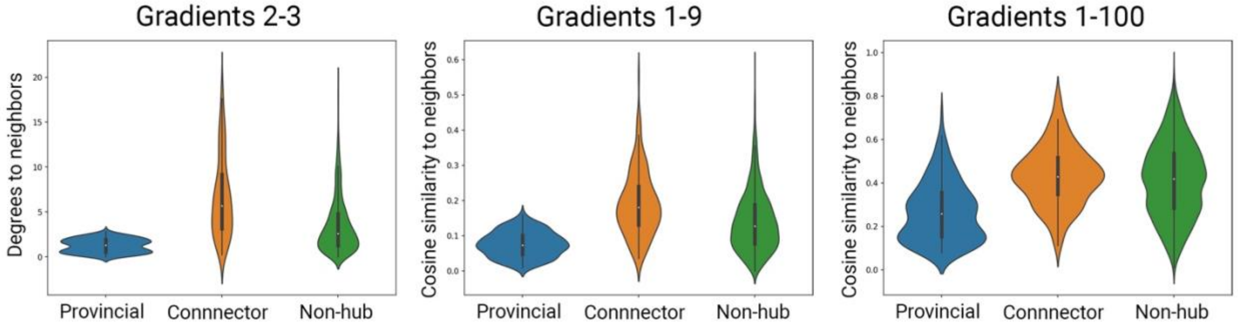


**Figure S8. Gradient similarity for hub regions and non-hubs.** Distributions of the similarity in gradient weights between regions and their most strongly functionally connected neighbors. The left plot shows the angular degrees to neighbors for provincial hubs, non-hubs, and connector hubs, analogous to **Figure 3C**. The middle and left plots showed the cosine distance of region gradient weights to neighbors when considering either latent space with either nine dimensions (middle) or 100 dimensions (right). In all cases, provincial hubs had significantly smaller angles to their neighbors than non-hubs, which in turn had smaller angles to their neighbors than connector hubs.

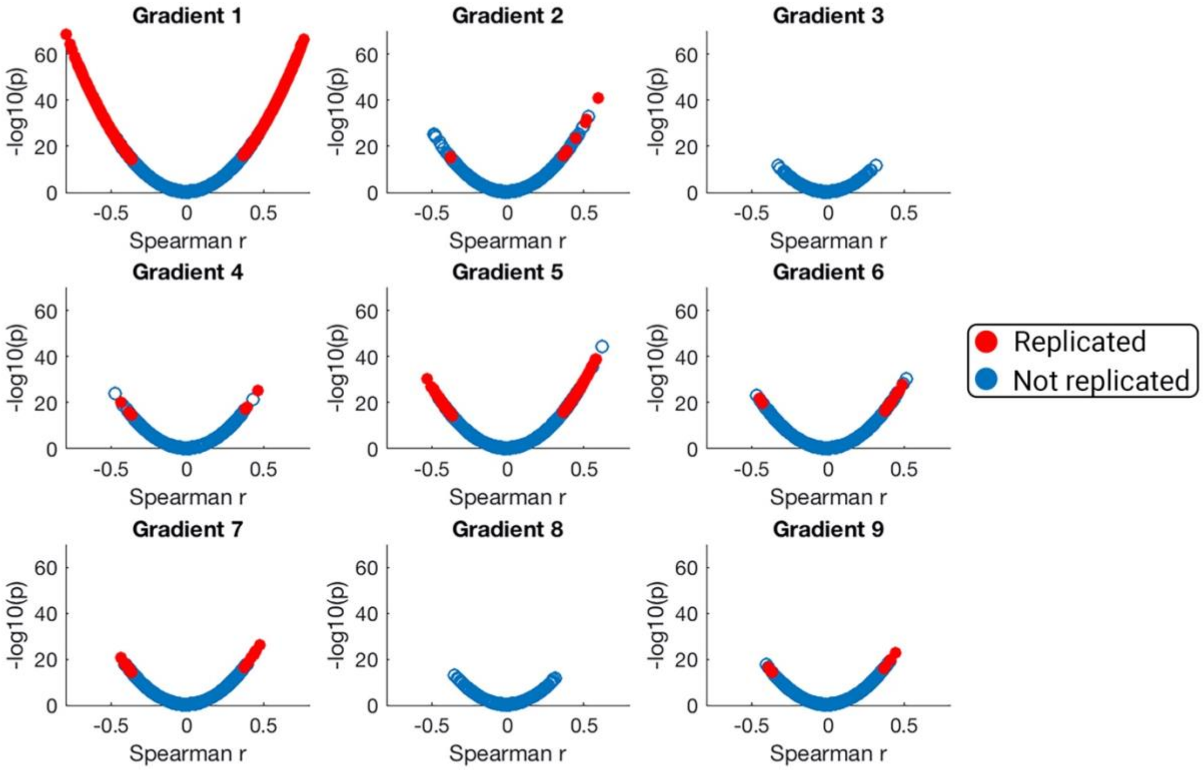


**Figure S9. Gradient/gene expression spatial correlations.** All 15655 correlation coefficients for the spatial correlation of 196-region gradient weights and 196-region gene expression levels are shown for gradients 1-9 from the discovery dataset, plotted against the corresponding log-transformed p-value. Genes that survived Bonferroni correction ($p < 3.55 \times 10^{-7}$) in both the discovery and validation datasets are shown in red.

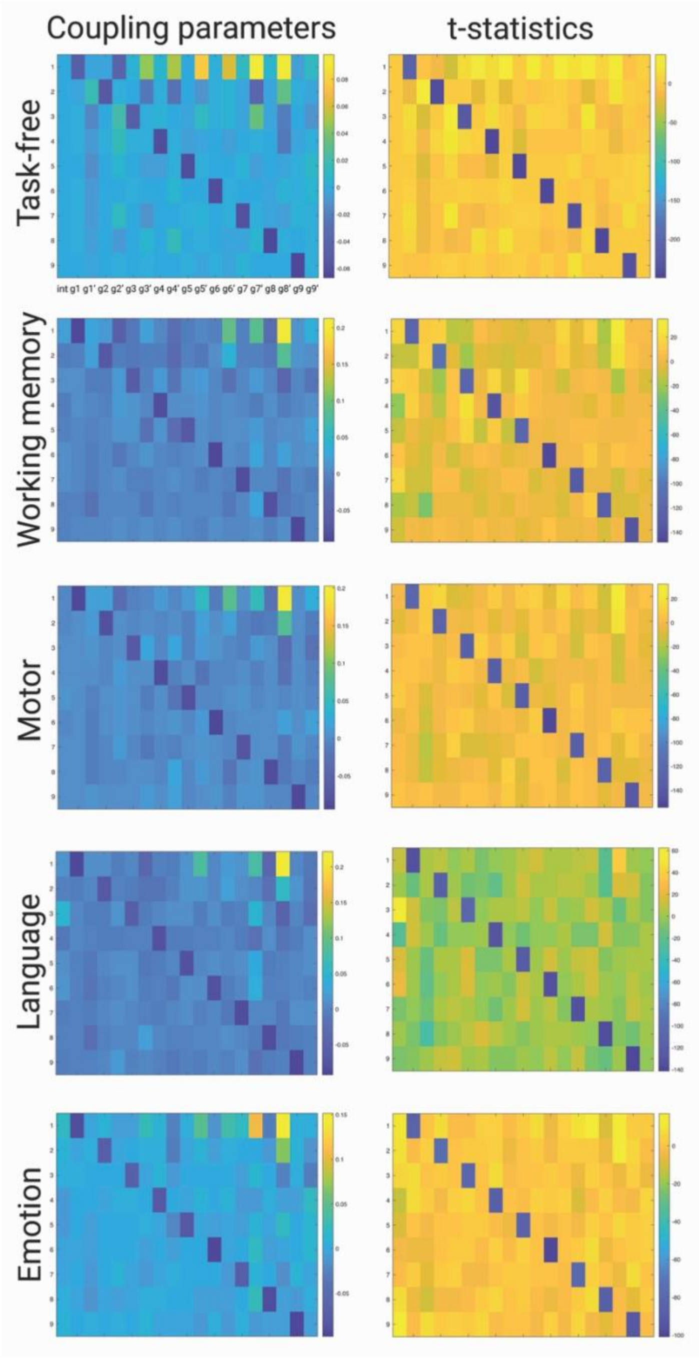


**Figure S10. Gradient coupling parameters.** The coupling parameters (beta values, left column) and associated t-statistics (right column) based on linear regression for each of the first nine gradients of gradient slope acceleration as a function of all gradients' slopes, slope velocities, and an intercept. Parameters are shown for the task-free state and the active condition of each task.

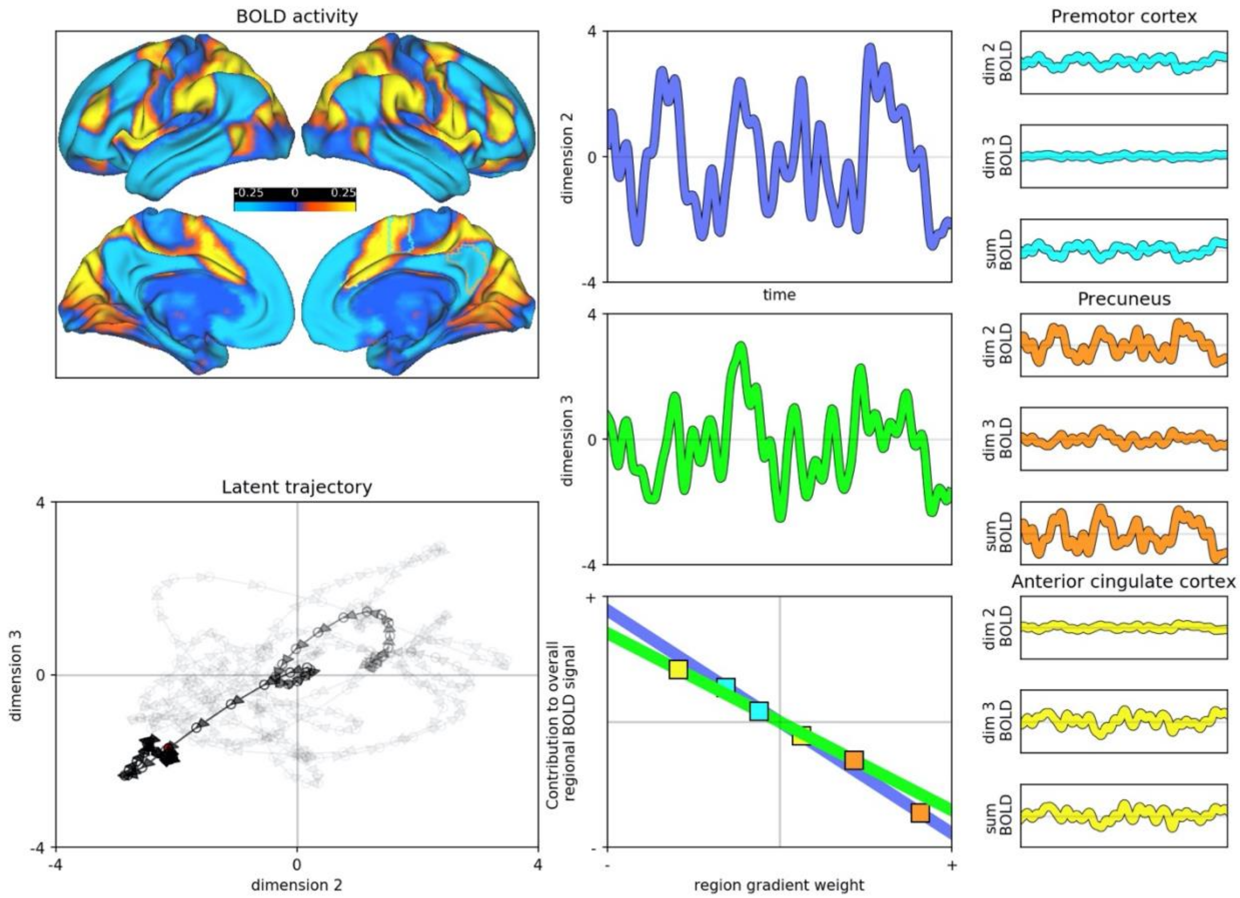


**Movie S1. Brain activity dynamics during task-free fMRI.** Activity is shown for one subject over 250 timepoints (3 minutes) of a task-free fMRI scan. The dynamic BOLD activity associated with latent dimensions 2 and 3 is shown (top left) along with the latent trajectory on dimensions 2 and 3 (bottom left), the gradient timeseries, and corresponding slopes (middle column). The regional BOLD activity timeseries are shown for three example regions as related to dimensions 2 and 3 and their summed result (right column).
